# Non-canonical IL-22 receptor signaling remodels the mucosal barrier during fungal immunosurveillance

**DOI:** 10.1101/2024.09.08.611873

**Authors:** Nicolas Millet, Jinendiran Sekar, Norma V. Solis, Antoine Millet, Felix E.Y. Aggor, Asia Wildeman, Michail S. Lionakis, Sarah L. Gaffen, Nicholas Jendzjowsky, Scott G. Filler, Marc Swidergall

**Author notes:** Correspondence: Marc Swidergall.

## Abstract

Mucosal barrier integrity is vital for homeostasis with commensal organisms while preventing pathogen invasion. We unexpectedly found that fungal-induced immunosurveillance enhances resistance to fungal outgrowth and tissue invasion by remodeling the oral mucosal epithelial barrier in mouse models of adult and neonatal *Candida albicans* colonization. Epithelial subset expansion and tissue remodeling were dependent on interleukin-22 (IL-22) and signal transducer and activator of transcription 3 (STAT3) signaling, through a non-canonical receptor complex composed of glycoprotein 130 (gp130) coupled with IL-22RA1 and IL-10RB. Immunosurveillance-induced epithelial remodeling was restricted to the oral mucosa, whereas barrier architecture was reset once fungal-specific immunity developed. Collectively, these findings identify fungal-induced transient mucosal remodeling as a critical determinant of resistance to mucosal fungal infection during early stages of microbial colonization.

## Main

Oral mucosal tissues are continuously exposed to food, airborne antigens, and commensal microbes, including fungi (*1*). The finely tuned implementation of innate and adaptive immune responses enables the host to maintain homeostasis with commensal species and neutralize invading organisms (*2-4*). As a central constituent of the mycobiome, the fungus *Candida albicans* colonizes the oral mucosa of up to 75% of healthy individuals (*5*). Settings of local or systemic immunosuppression result in oropharyngeal candidiasis (OPC) (*6*). Since, in healthy individuals *C. albicans* causes no harm, fungal colonization appears to be evolutionarily selected for appropriate metabolic function and immune priming (*7-10*). While *C. albicans* has been extensively studied as a pathogen (*11-14*), the primary lifestyle of this fungus in the oral cavity is as a commensal (*15-17*). The natural diversity in *C. albicans* influences the outcome of the interaction between the fungus and the host (*16*) implying that the involvement of specific immune pathways to the host defense against *C. albicans* is modulated in a strain-dependent manner. In mouse models of OPC, *C. albicans* strains can be grouped into two categories: pathogenic-like (PL) and commensal-like (CL) (*16*). While PL *C. albicans* strains induce a strong pro-inflammatory response that leads to rapid clearance in the oral mucosa (acute OPC), CL *C. albicans* strains instead trigger a tempered inflammatory response that permits long-term fungal colonization, thus mimicking commensal colonization in humans. Still, the regulation of mucosal homeostasis during commensal fungal colonization remains poorly understood.

The epithelial architecture of mucosal surfaces such as the oral cavity is crucial for its host defensive function (*18-20*), providing structural immunity by initiating and coordinating immune responses (*21*). However, relatively little is known about how this is coordinated in the oral mucosa. Furthermore, the epithelium balances a multiplicity of roles in early life (*22*), while the acquired microbiome contributes to the development of immunity in newborns. Following exposure, the mucosal immune system of neonates undergoes successive, non-redundant phases that support the developmental needs of the infant to establish immune homeostasis (*23*). While tissue remodeling has been associated with pathological features post-injury or disease (*24, 25*), commensal organisms may induce changes in barrier structures to promote homeostasis (*26*).

Here, we studied mucosal immune responses and tissue homeostasis in mouse models of persistent *Candida* oral colonization at distinct ranges of age. We show that immunosurveillance-induced epithelial expansion and remodeling mediates resistance against fungal outgrowth during the onset of fungal colonization. IL-22, a critical cytokine in epithelial homeostasis and host defense at mucosal surfaces, has long been associated with signaling through a well-characterized receptor complex (*27*). Traditionally, IL-22 exerts its protective effects trough binding to the IL22RA1-IL10RB receptor complex. However, our study reveals that IL-22 recognition and signaling extend beyond this canonical pathway, expanding the current understanding of its biological roles. Oral mucosal remodeling required IL-22-mediated gp130 activation in non-canonical cytokine receptor complexes with IL-22RA1 and IL-10RB. IL-22 mediated oral epithelial remodeling was transitory and a subsequent mucosal remodeling event in later stages of colonization required *Candida*-specific immunity. Finally, fungal-induced mucosal remodeling prevents tissue invasion in a mouse model of neonatal colonization. These findings provide novel insights into the molecular mechanisms of IL-22-mediated signaling, constituting a new pathway for antifungal immunity, and expand our understanding of microbe-induced epithelial remodeling and homeostasis trough a non-canonical receptor complex.

### Persistent *C. albicans* colonization induces epithelial remodeling

In a mouse model of OPC (*28*), acute infection with a prototypic PL *C. albicans* strain (SC5314) leads to its rapid clearance from the oral cavity, whereas a CL *Candida* strain (CA101) persists int he oral mucosa (Fig. S1)(*15*). Several host cell types respond to *C. albicans* encounter in the oral cavity. To develop a single cell transcriptome profile of the oral mucosa, we evaluated gene expression in tissues of CL colonized mice and from mice after pathogenic *Candida* clearance (Fig. 1A). Our analysis identified 18 distinct cell subpopulations, which were present in both, CL colonized mucosa and mucosal tissue after PL clearance (Fig. 1B). The identified subpopulations expressed cell-type specific marker genes (Fig. S2) consistent with classical, well-established markers for each respective cell population. The scRNA-sequencing analysis revealed that the epithelial proportions increased during persistent fungal colonization (Figure 1C). Accordingly, epithelial cells from colonized mice had higher expression of proliferation marker genes (*Mki67, Cenpf, Cenpa*) (Fig. 1D). Tissue histology during the onset of CL *Candida* colonization revealed that the epithelial layer expanded (Fig. 1E) and basal epithelial cell proliferation increased indicated by Ki67 staining (Fig. 1F, G). Next, we performed RNA sequencing of epithelial-enriched mucosal tissue (Fig. 1H). Using Gene Set Enrichment Analysis (GSEA), we found that genes involved in keratinization and keratinocyte differentiation were significantly enriched in fungal colonized epithelial tissues (Fig. 1I). Similarly, genes for keratinization were enriched in epithelial cells from colonized mice in the scRNA-sequencing data set (Fig. S3). Keratins influence the epithelial architecture by determining cell compartmentalization and differentiation (*29*). Several keratins, including keratin 14 (K14), were highly expressed in the mucosa of colonized mice (Fig. 1J). Basal epithelial cells express K14 in all regions, while the suprabasal epithelial layer (SEL) expresses K13 (*30*). Consistent with these findings the K14 epithelial layer expanded during commensal colonization, while the epithelium had similar architecture after PL encounter and clearance compared to Sham infection (Fig.1K, L). Notably, the K13 layer distribution and thickness remained unchanged during CL colonization (Fig. S4). Time course studies revealed that K14 epithelial expansion occurs after 8 days of fungal colonization (Fig. 1M, S5). Collectively, these data show that persistent fungal colonization induces distinct epithelial subset expansion; thus, indicating that *Candida* commensalism influences the oral mucosal architecture.

**Figure 1.**
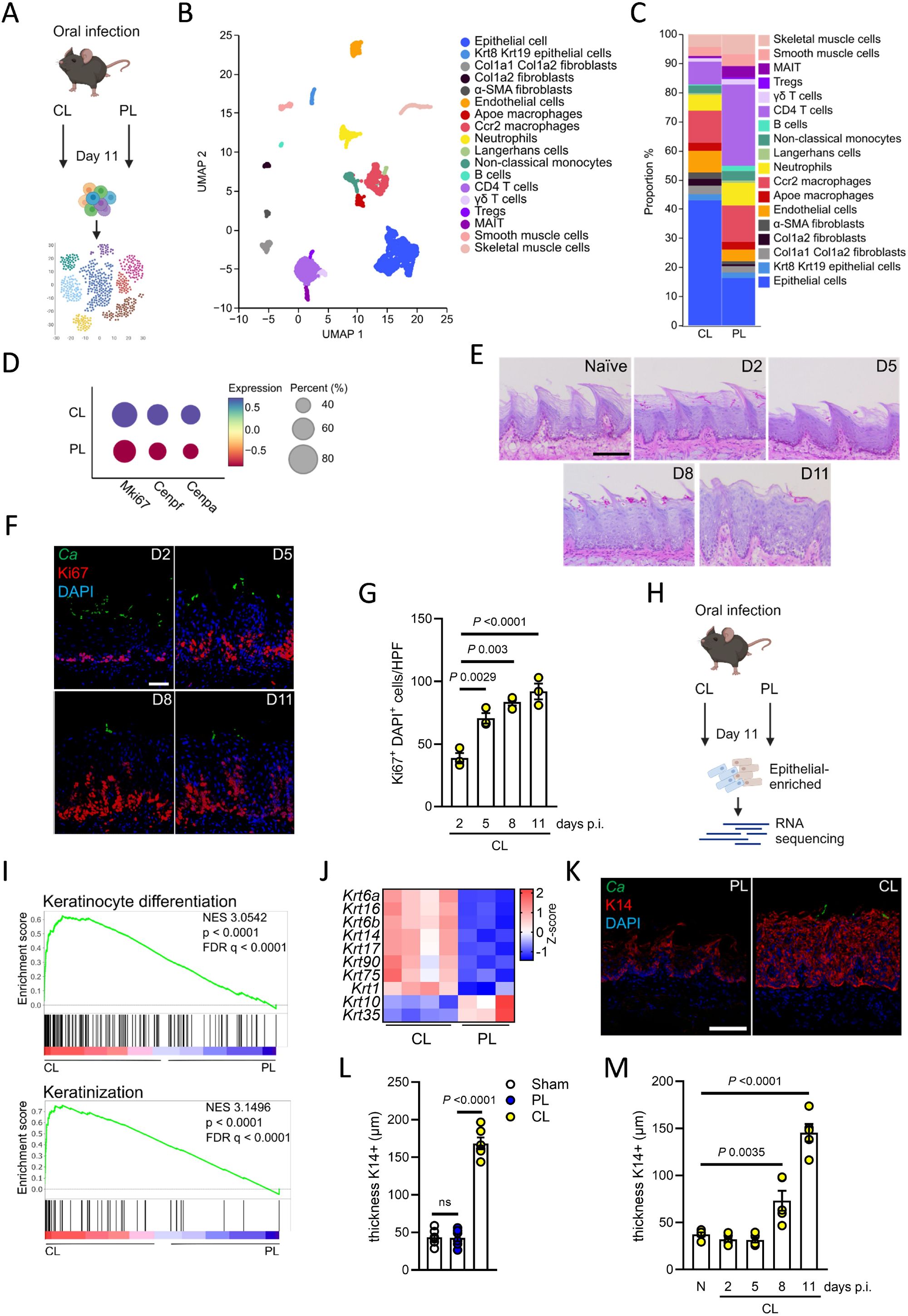
Oral *C. albicans* colonization induces epithelial remodeling. **A** Experimental setup for single cell collection and sequencing. **B** Cell types identified in the oral mucosa after 11 days of CL and PL infection with UMAP projections of scRNA-seq data. **C** Proportions of identified cell types separated by PL and CL infection. **D** Expression of *Miki67, Cenpf*, and *Cenpa* in epithelial cells of PL and CL infected mice. **E** Representative pictures of PAS staining of CL-infected mice over time. **F** Representative pictures of Ki67 staining of CL-infected mice over time. Scale bar 50μm. **G** Quantification of Ki67^+^ DAPI^+^ cells CL-infected mice over time. *N* = 3; Ordinary one-way ANOVA. **H** Experimental setup for sequencing of epithelial-enriched tissues. **I** GSEA of keratinocyte differentiation and keratinization sequencing data of epithelial-enriched tissues. *N* = 3-4. **J** Z-scores of keratin genes in epithelial-enriched tissues after PL and CL infection (Day 11). *N* = 3-4 **K** Representative pictures of K14 staining of PL- and CL-infected mice after 11 days. **L** Quantification of K14 thickness in Sham-, PL, and CL-infected mice after 11 days. *N* = 6; combined data of two independent experiments. Ordinary one-way ANOVA. **M** Quantification of K14 thickness in CL-infected mice over time. *N* = 5; combined data of two independent experiments. Ordinary one-way ANOVA. PL, pathogen-like; CL, commensal-like.

### CD4 T cells are a major source of IL-22 during commensal colonization

T cells play vital roles in the mucosal antifungal immunity (*31, 32*), while some CD4^+^ T cells reside in the oral mucosa of healthy individuals (*20*). In fact, T cells can promote stromal cell proliferation through secretion of cytokines (*33-35*). Thus, we compared protein expression kinetics of various T helper (Th) cell-associated cytokines and chemokines during persistent *C. albicans* colonization and acute OPC. Acute infection with the PL *C. albicans* strain led to early TNFα and IL-1β induction, whereas the CL *Candida* strain colonized the oral mucosa without inducing inflammation (Fig. 2A). In acute OPC, IL-17 and IL-22 are expressed by Type 17 cells with similar kinetics (*11, 36-38*). In contrast, our data show that IL-22 and IL-17 are differentially expressed during persistent colonization (Fig. 2A, B). Mice infected with a PL strain induced IL-17A and IL-22 similarly followed by rapid decline after fungal clearance. However, in CL colonized mice, mucosal IL-22 levels remained high, while IL-17A levels were low at the onset of colonization (Fig. 2B). IL-22 neutralization after CL *C. albicans* colonization (Fig. 2C) resulted in fungal proliferation (Fig. 2D), while antibody neutralization of IL-17A did not alter the fungal burden in colonized mice. This suggested that IL-22 may have a prominent role in controlling fungal outgrowth after mucosal colonization. Next, we determined the cellular origin of IL-22. Our scRNA-sequencing dataset suggested that CD4 T cells highly express *Il22* during persistent fungal colonization (Fig. 2E). Using *IL22TdTomato* reporter mice, we confirmed that CD4 T cells are the major source of IL-22 during colonization with CL *Candida* (Fig. 2F). Next, we evaluated localization of CD4 T cells during commensal colonization within the oral mucosa. CD4 T cells were exclusively found in the epithelial and submucosal layers of commensal colonized mice, which differed markedly from PL- or Sham-infected mice (Fig. 2G). Within the helper T cells, IL-22 is mainly produced by Th17 and Th22 subsets (*39*). *Ex vivo* stimulation of mucosal resident CD4 T cells from commensal colonized and pathogenic infected mice showed that during commensal colonization Th17 and Th22 cells are the major source of IL-22, while Th22 cell frequencies increased (Fig. 2H, I).

**Figure 2.**
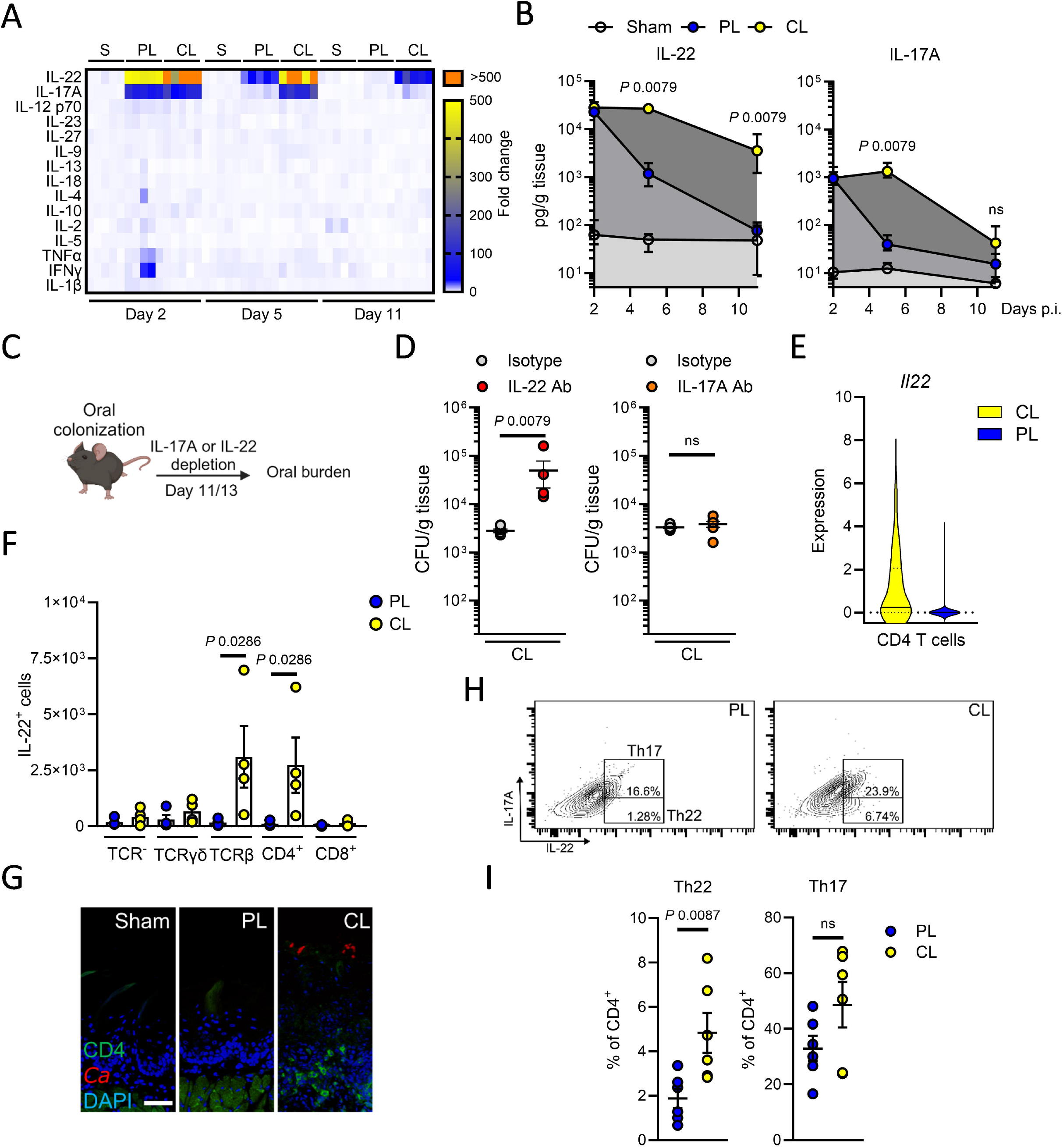
Th17/Th22 cells produce IL-22 during *C. albicans* colonization. **A** Heat map presented in fold change of various cytokines during PL, CL, and Sham infection over time *N* = 10; combined data of two independent experiments. Two-tailed Mann–Whitney Test. **B** Levels of IL-17A and IL-22 in PL-, CL-, and Sham-infected mice; *N* = 10. Two-tailed Mann–Whitney Test. **C** Scheme of IL-17 and IL-22 depletion during CL-colonization. **D** Oral fungal burden of mice colonized with CL after treatment on day 11 and 13 with anti-IL-17A or anti-IL-22 antibodies. *N* = 6; combined data of two independent experiments. Two-tailed Mann–Whitney Test. The y-axis is set at the limit of detection (20 CFU/g tissue). **E** Expression of *Il22* in CD4 T cells identified in the single cell RNA-sequencing data set. **F** Quantification of IL-22 expressing cells during CL colonization using *IL22TdTomato* reporter mice. *N*=4. Two-tailed Mann–Whitney Test. **G** Representative immunofluorescence pictures of localization of CD4 T cells in tissues after 11 days of infection. Scale bar 50μm. **H** Representative flow cytometry plots for intracellular IL-17A and IL-22 levels in CD4+ cells. **I** Levels of Th17 and Th22 cells after 11 days of infection. *N* = 6; combined data of two independent experiments. Two-tailed Mann–Whitney Test. PL, pathogen-like; CL, commensal-like.

### Epithelial expansion depends on IL-22 signaling via non-canonical gp130 receptor complexes

The IL-17- and the IL-22 receptors, which mediate anti-*C. albicans* immunity during acute OPC, play distinct and restricted roles in distinct sublayers of the stratified oral epithelium (*36*). During CL-*Candida* colonization, IL-22RA1 protein expression extended into the suprabasal layer (Fig. S6) consistent with K14 expression (Fig.1K). IL-22 has been implicated in multiple aspects of epithelial barrier function and wound repair, including regulation of cell growth (*40*). Accordingly, IL-22 induced proliferation of human oral epithelial cells in a dose-dependent manner, while high cytokine concentrations inhibited oral epithelial cell growth (Fig. 3A, S7). Similarly, IL-22 deficient mice and IL-22 depletion during CL colonization reduced K14 epithelial layer expansion and proliferation (Fig. 3B-D, Fig, S8), as well as resistance against fungal outgrowth (Fig. 3E) in the setting of intact IL-17 signaling (Fig. S9). IL-22-mediated proliferation required JAK-TYK2-STAT3 signaling in human oral epithelial cells (Fig. S10). Classical IL-22 signal transduction is mediated by binding of the cytokine to a receptor complex consisting of IL-22RA1 and IL-10RB (*27*). However, while mice deficient for either receptor, IL-22RA1 and IL-10RB respectively, were more susceptible to CL *Candida* outgrowth (Fig. 3F), the K14 layer surprisingly still underwent expansion in the absence of either receptor chain (Fig. 3G, S11). Notably, IL-10 signals through a heterotetrameric complex comprising of IL10Rα and IL10RB but was dispensable for fungal control and epithelial remodeling (Fig. S12). Recently, studies in synthetic cytokine biology have challenged the classical view of IL-22 signaling via a heterodimeric IL-22RA1 and IL-10RB complex. Synthetic cytokine receptor chains of IL-22RA1 and IL-10RB form functional heterodimeric receptor signaling complexes with a IL-6 receptor chain of gp130 to induce STAT3 signal transduction (*41*). Proximity ligation assays and co-immunoprecipitation identified the classical heterodimeric IL-22RA1 and IL-10RB complex, while revealing that IL-10RB, as well as IL-22RA1, form receptor complexes with gp130 in human oral epithelial cells (Fig. 3H, S13). We tested whether antibody blockade of either receptor chain, IL22RA1 and IL-10RB, would abolish STAT3 signal transduction in human oral epithelial cells. While antibodies blocking IL-22RA1 or IL-10RB induced similar STAT3 activation following IL-22 exposure, treatment with both antibodies reduced epithelial activation and IL-22 mediated proliferation (Fig. 3I, J, S14). Next, we determined the contribution of gp130 signaling to IL-22-mediated oral epithelial proliferation. Pharmacological inhibition of gp130 signaling reduced IL-22-mediated oral epithelial proliferation *in vitro* (Fig. 3K), while reducing STAT3 activation (Fig. 3L, S14). IL-6 binds to membrane-bound IL-6R to induce signaling via gp130 (*42*). However, IL-22-mediated epithelial proliferation was independent of IL-6-gp130 signaling (Fig. S15). Of note, STAT3 activity remained unchanged during inhibition of gp130 in intestinal epithelial cells (Fig. S16), suggestive of organ-specific receptor complex signaling in response to IL-22. Inhibition of gp130 signaling during oral CL persistence increased the fungal burden and inhibited K14 expansion and remodeling (Fig. 3M-P). Thus, IL-22 signals via various receptor chain combinations to provide oral mucosal antifungal immunity during fungal immunosurveillance, while oral epithelial proliferation and expansion depends on non-canonical gp130 signaling receptor complexes.

**Figure 3.**
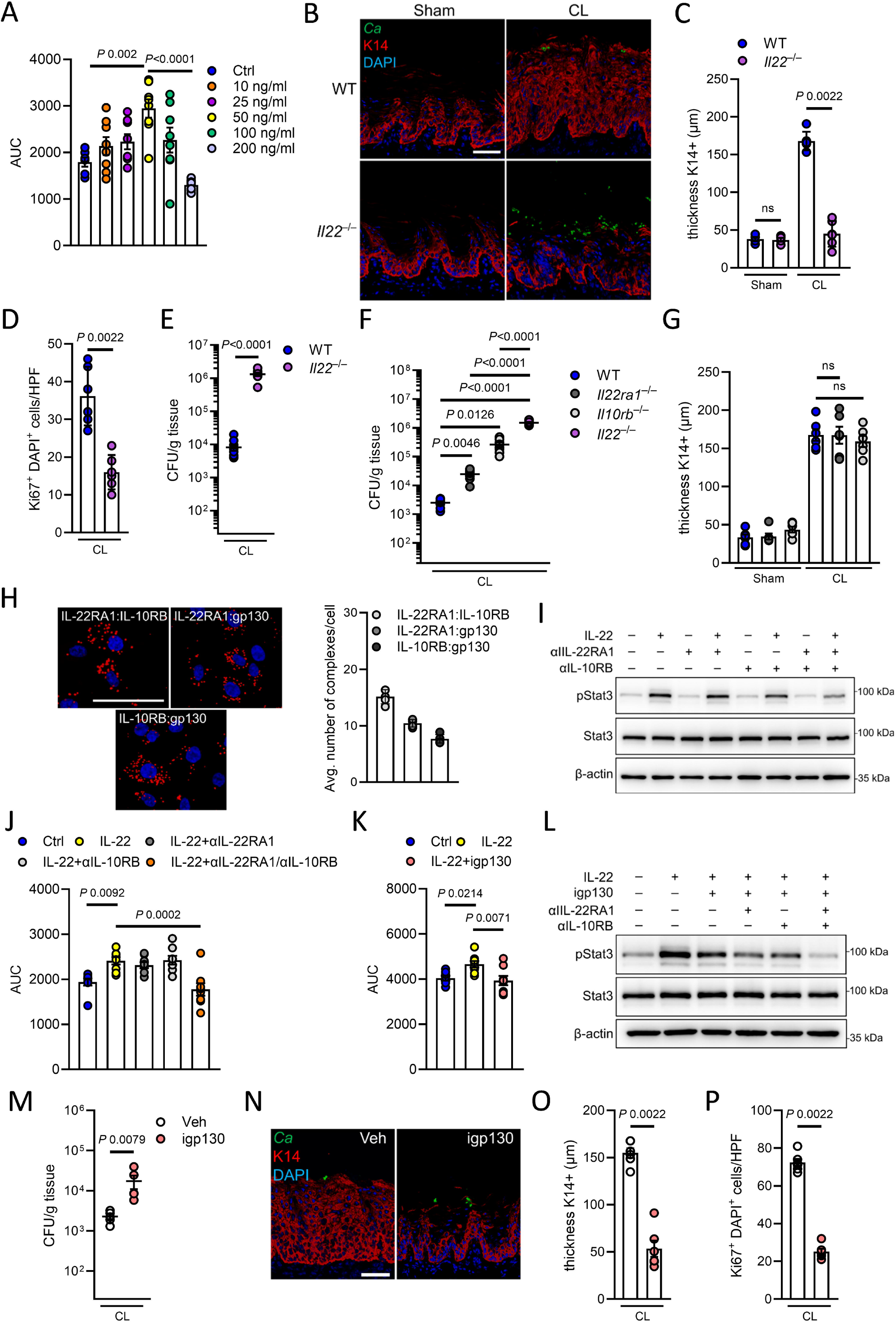
Epithelial expansion requires non-classical IL-22 signaling. **A** Area under the curve (AUC) of oral epithelial cells in response to different concentrations of IL-22. Growth was determined by confluence over time. N=8. Ordinary one-way ANOVA. **B** Representative pictures of K14 staining of Sham- and CL-infected mice after 11 days. Scale bar 50μm. **C** Quantification of K14 thickness in Sham- and CL-infected WT and *Il22*^−/–^ mice at day 11. *N* = 6; combined data of two independent experiments. Two-tailed Mann–Whitney Test. **D** Quantification of Ki67^+^ DAPI^+^ cells CL-infected mice over time. *N* = 6; combined data of two independent experiments. Two-tailed Mann–Whitney Test. **E** Oral fungal burden of WT and *Il22*^−/–^ mice colonized with CL after 11 days. *N* = 8-9; combined data of two independent experiments. Two-tailed Mann–Whitney Test. The y-axis is set at the limit of detection (20 CFU/g tissue). **F** Oral fungal burden of indicated mice colonized with CL after 11 days. *N* = 7; combined data of two independent experiments. Brown-Forsythe and Welch ANOVA. The y-axis is set at the limit of detection (20 CFU/g tissue). **G** Quantification of K14 thickness in indicated mice at day 11. *N* = 6; combined data of two independent experiments. Ordinary one-way ANOVA. **H** (Left) Representative Images for proximity ligation assay (PLA) for indicated receptor complexes. Scale bar 50μm. Red dots indicate receptor complexes. (Right) Quantification of average number of complexes per cell. *N* = 5. **I** Representative immunoblot of STAT3 activation during IL-22 incubation in the presence of IL-22RA1, IL-10RB, or combination of IL-22RA1/IL-10RB blocking antibodies. **J** Area under the curve (AUC) of oral epithelial cells in response to IL-22 treatment in the presence of IL-22RA1, IL-10RB, or combination of IL-22RA1/IL-10RB blocking antibodies. Growth was determined by confluence over time. *N* = 8. Ordinary one-way ANOVA. **K** Area under the curve (AUC) of oral epithelial cells in response to IL-22 treatment in the presence of the gp130 inhibitor SC144 (igp130). Growth was determined by confluence over time. *N* = 8. Ordinary one-way ANOVA. **L** Representative immunoblot of STAT3 activation during IL-22 incubation in the presence of gp130 inhibitor, IL-22RA1, IL-10RB, or combination. **M** Oral fungal burden of mice treated with gp130 inhibitor (igp130) or vehicle control (Veh) colonized with CL after 11 days. *N* = 6; combined data of two independent experiments. Two-tailed Mann–Whitney Test. The y-axis is set at the limit of detection (20 CFU/g tissue). **N** Representative pictures of K14 staining of Veh- and igp130 treated mice after 11 days of CL infection. Scale bar 50μm **O** Quantification of K14 thickness in Veh- and igp130 treated mice after 11 days of CL infection. *N* = 6; combined data of two independent experiments. Two-tailed Mann–Whitney Test. **P** Quantification of Ki67^+^ DAPI^+^ cells CL-infected mice in the presence and absence of gp130 signaling. *N* = 6; Two-tailed Mann–Whitney Test. PL, pathogen-like; CL, commensal-like.

### Transient epithelial expansion depends on development of *Candida*-specific immunity

While oral CL infection leads to epithelial expansion at the onset of colonization, at later time points of fungal persistence the K14 expansion is reset to levels comparable to the baseline cell turnover seen in naïve mice (Fig. 4A-C), suggesting that mucosal remodeling is transitory, regulated, and depends on an auxiliary mechanism. A unifying theme of susceptibility to mucocutaneous candidiasis is seen in both humans and mice with a variety of genetic defects within the IL-17 pathway (*3, 31, 43*). The generation of organism-specific adaptive immune responses takes time to generate sufficient cells, due to the inherent demands for extensive proliferation and differentiation of naive cells into effector cells (*44*). Thus, we tested if *Candida*-specific immunity is required to remodel the oral mucosal barrier in subsequent phases of colonization. During late stages of fungal persistence, increased numbers of *Candida*-specific IL-17 secreting cells in cervical lymph nodes (Fig. 4D) inversely correlated with reversion of the K14 barrier thickness (Fig. 4E) suggesting the requirement of *Candida*-specific immunity to reset barrier architecture. *Rag1*^−/–^ were more susceptible to fungal outgrowth at late stages during colonization (Fig. 4F), while K14 epithelial expansion remained high (Fig. 4G, H). Collectively, we show that IL-22-mediated oral epithelial remodeling is transitory and requires *Candida*-specific immunity for a subsequent mucosal remodeling event in later stages of colonization.

**Figure 4.**
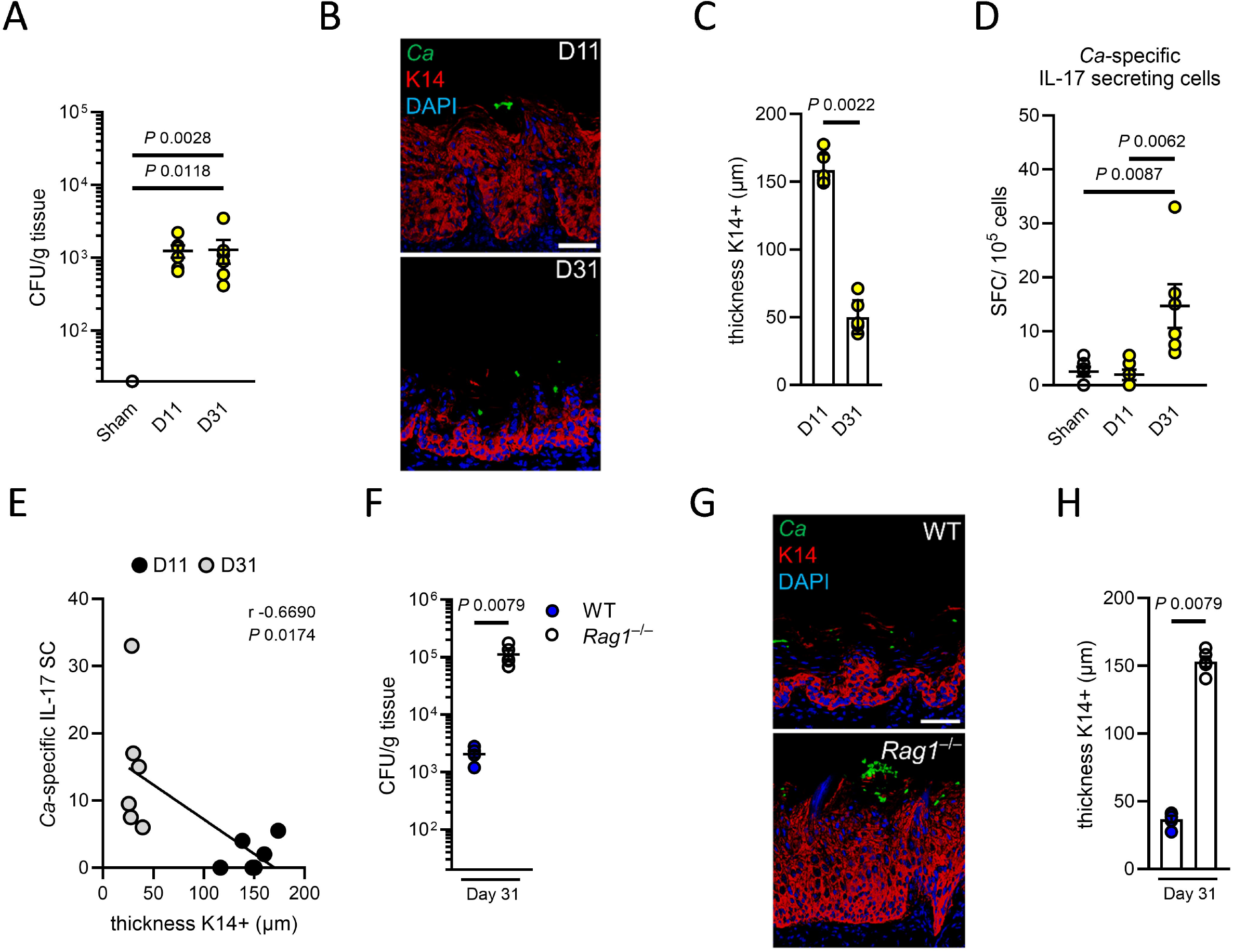
Transitory epithelial K14 expansion depends on *Candida*-specific immunity. **A** Oral fungal burden of mice colonized with CL after indicated time points. *N* = 6; combined data of two independent experiments. Two-tailed Mann–Whitney Test. The y-axis is set at the limit of detection (20 CFU/g tissue). **B** Representative pictures of K14 staining of CL-infected mice after 11 and 31 days. Scale bar 50μm. **C** Quantification of K14 thickness in CL-infected mice at indicated time points. *N* = 6; combined data of two independent experiments. Two-tailed Mann– Whitney Test. **D** Determination of IL-17 secreting cells isolated from cervical lymph nodes mice and stimulated with *Candida* antigen pools for 24 hours using ELISpot. N=6. Ordinary one-way ANOVA. **E** Correlation of K14 thickness and *Candida* (*Ca*)-specific IL-17 secreting cells. N=6. Correlation was determined by Pearson. **F** Oral fungal burden of WT and *Rag1*^−/–^ mice colonized with CL after 31 days. N=5; combined data of two independent experiments. Two-tailed Mann– Whitney Test. The y-axis is set at the limit of detection (20 CFU/g tissue). **G** Representative pictures of K14 staining of CL-infected mice after 31 days. Scale bar 50μm. **H** Quantification of K14 thickness in CL-infected mice after 31 days. *N* = 5; combined data of two independent experiments. Two-tailed Mann–Whitney Test. PL, pathogen-like; CL, commensal-like.

### Commensal-induced epithelial expansion prevents fungal tissue invasion in neonates

Neonates exhibit differentially adapted immune responses when compared to adults (*45*), with important implications for immunity to OPC; indeed, oral thrush is commonly seen in infants (*46*), though the underlying basis for this is not well defined. In this regard, neonatal umbilical cord blood-derived CD4 T cells are intrinsically less able to differentiate into Th17 cells, but rather tend to skew towards an IL-22 phenotype (*47*). Here, we established a mouse model of neonatal fungal colonization to study early-life interactions between the oral epithelium and *Candida*, which occurs in humans (Fig. 5A, B). Consistent with our findings in adult animals, oral CL colonization in neonates induced K14 epithelial expansion (Fig. 5C, D) and CD4 T cell infiltration at the onset of colonization (Fig. S17). Neonatal mice deficient in IL-22 had increased oral fungal burden, abrogated K14 expansion, and decreased fungal-induced epithelial proliferation (Fig. 5F-H). Furthermore, IL-22 deficiency in neonates increased the depth of fungal tissue invasion (Fig. 5I, J) which was not observed in adult *Il22*^−/–^ mice (Fig. 5J) suggesting that IL-22-mediated immunity and associated tissue remodeling plays a critical role preventing fungal invasion during early life, highlighting the importance of IL-22 in neonatal immune defense and epithelial protection.

**Figure 5.**
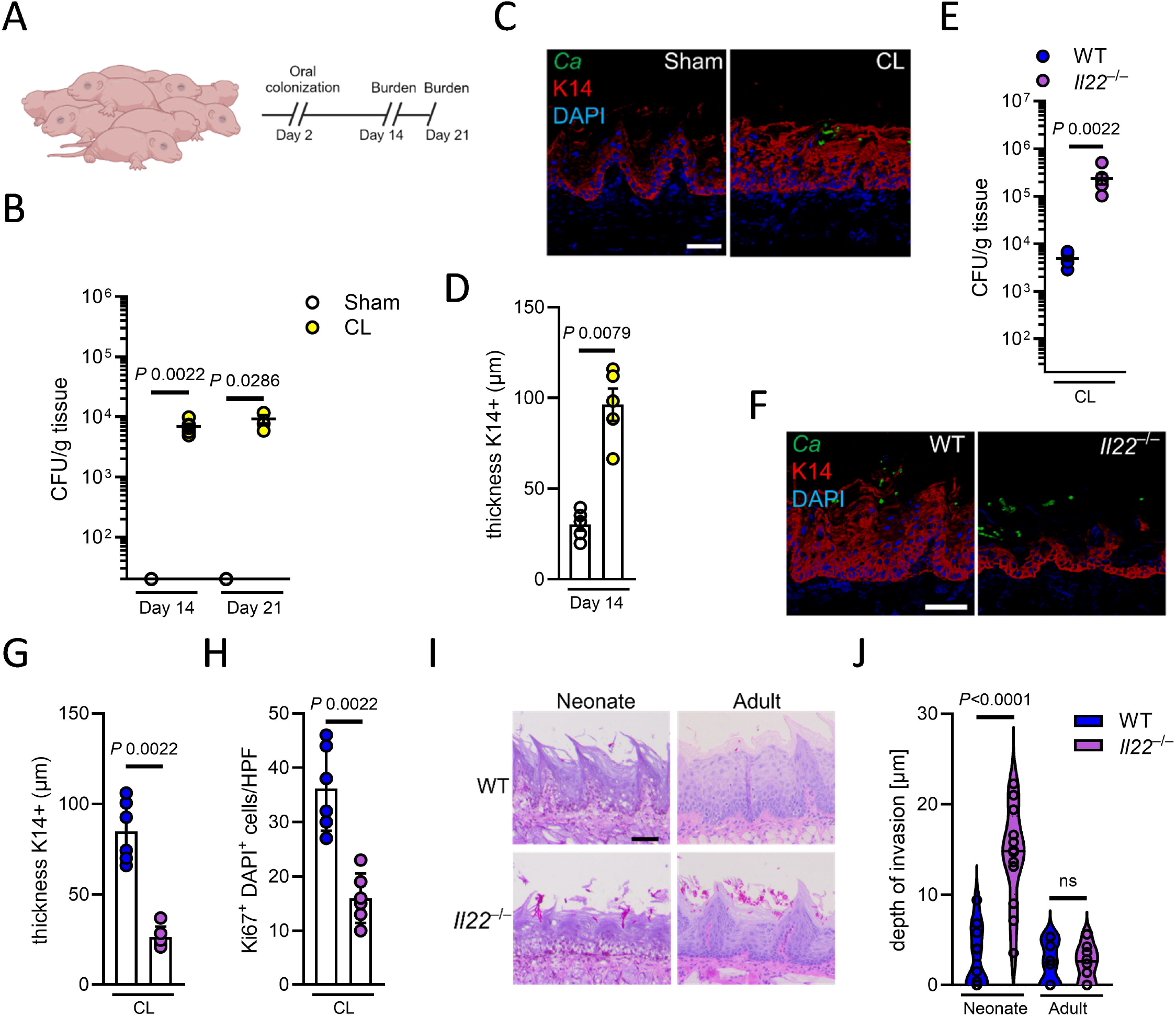
Epithelial remodeling prevents fungal invasion during neonatal colonization. **A** Experimental setup for neonatal colonization of wild type mice. **B** Oral fungal burden of neonatal mice Sham-infected or colonized with CL after indicated time points. *N* = 4-6; combined data of two independent experiments. Two-tailed Mann–Whitney Test. The y-axis is set at the limit of detection (20 CFU/g tissue). **C** Representative pictures of K14 staining of Sham-infected or CL-infected mice at day 14 of life. Scale bar 50μm. **D** Quantification of K14 thickness in Sham-infected or CL-infected mice at day 14 of life. *N* = 5; combined data of two independent experiments. Two-tailed Mann–Whitney Test. **E** Oral fungal burden of neonatal WT and *Il22*^−/–^ mice colonized with CL at day 14 of life. *N* = 6; combined data of two independent experiments. Two-tailed Mann– Whitney Test. The y-axis is set at the limit of detection (20 CFU/g tissue). **F** Representative pictures of K14 staining of WT and *Il22*^−/–^ mice colonized with CL at day 14 of life. Scale bar 50μm. **G** Quantification of K14 thickness in neonatal WT and *Il22*^−/–^ mice colonized with CL at day 14 of life. *N* = 5; combined data of two independent experiments. Two-tailed Mann–Whitney Test. **H** Quantification of Ki67^+^ DAPI^+^ cells in neonatal WT and *Il22*^−/–^ mice colonized with CL at day 14 of life. *N* = 6; combined data of two independent experiments. Two-tailed Mann–Whitney Test. **I** Representative PAS staining of tongue tissues of neonatal (day 14 of life/day 12 of CL colonization) and adult (day 11 of CL colonization) WT and *Il22*^−/–^ mice. Scale bar 50μm. **J** Quantification of depth of fungal invasion in neonatal WT and *Il22*^−/–^ mice colonized with CL at day 14 of life and adult mice after 11 days of CL colonization. *N* = 8-12; combined data of four independent experiments. Two-tailed Mann–Whitney Test. PL, pathogen-like; CL, commensal-like.

## Discussion

Resistance to mucosal fungal infection is tightly linked to maintenance of barrier integrity (*20, 48*). The communication between epithelial and immune cells enables coordinated responses that maintain homeostasis and elicit host defenses (*21, 49*). Our findings support a paradigm by which fungal-induced immunosurveillance in the oral mucosa leads to transitory mucosal tissue remodeling and enhanced barrier resistance to fungal outgrowth.

The microbiome is a crucial factor for shaping and modulating immune system responses through cytokine signatures (*50*). As such, our data unveil an IL-22-mediated pathway that critically regulates mucosal antifungal host defense in mice following continued exposure to fungi. We demonstrate that oral epithelial IL-22 signaling relies on various receptor chain combinations to mediate antifungal immunity, while gp130 receptor complexes drive epithelial remodeling and proliferation, which expands on reported tissue-specific protective roles of IL-22 receptor signaling (*51*). Our findings of a non-canonical IL-22 receptor complex to orchestrate epithelial remodeling represents a novel mechanism of mucosal antifungal defense, challenging the classical understanding of IL-22 signaling. The IL-22 recognizing receptor complexes contribute to different degrees to oral antifungal immunity indicated by diverse level of fungal dysbiosis. These findings align with clinical phenotypes of inborn errors of immunity in IL-10RB, in which patients are resistant to oral candidiasis (*52*) while a small fraction of patients with gp130-dependent Hyper-IgE syndrome develop chronic mucocutaneous candidiasis (*53, 54*). Future studies will be required to determine additive and compensatory mechanisms of the IL-22RA1, IL-10RB, and gp130 epithelial receptor complexes during fungal colonization and infection in the oral mucosa. Inhibition of gp130, as well as dual receptor blockage of IL-22RA1 and IL-10RB, impairs IL-22-mediated STAT3 phosphorylation and epithelial proliferation, while some transcription factor activity remains. T cell differentiation requires optimal STAT3 phosphorylation (*55*) suggesting a similar mechanism requiring fine-tuned STAT3 activity exist in epithelial cells to induce proliferation. On the other hand, IL-22 signaling via IL-22RA1, IL-10RB, and gp130 complexes is organ- and context dependent since IL-22-mediated STAT3 activation in intestinal epithelial cells occurs independent of gp130.

*Candida*-responsive CD4^+^ T cells are primed in all healthy individuals as a consequence of exposure to the commensal fungus (*4*). Predominantly characterized by a Th17 profile, *C. albicans*-specific T cells are detected in epithelial tissues at the onset of colonization in experimental models of mucosal infection (*56, 57*). During the initial phases of acute mucosal infection in a naïve host, anti-*Candida* immunity is driven by epithelial pattern recognition and damage, and subsequent innate and bystander T cell responses (*13, 58*). Within the oral mucosa, tissue-resident Flt3L-dependent dendritic cells (DCs) and CCR2-dependent monocyte-derived DCs collaborate in fungal antigen presentation and T cell priming (*59*). The generation of antigen-specific immunity depends on its nature and the site of exposure. Transitory epithelial proliferation and oral mucosal remodeling enhances resistance to fungal outgrowth, while reversal of the epithelial expansion occurs once *Candida*-specific immunity is developed (Fig. S18). The sustained epithelial expansion and increased fungal burden in mice lacking fungal-specific immunity underscore the importance of functional adaptive immunity in controlling both, epithelial remodeling and fungal persistence.

During early life, newborns encounter an abundance of antigenic challenges derived from commensal and pathogenic organism. Although 5–7% of infants develop oral candidiasis (*46*), aberrant cellular innate immune responses and an inexperienced adaptive immune system do not elaborate high resistance against fungal outgrowth. Thus, neonates may exhibit a degree of immunological tolerance to fungal colonization to prevent harmful immune reactions in mucosal tissues. After birth, the neonatal epithelium is permeable, while lacking pattern recognition receptors (*60, 61*) suggesting that neonatal oral epithelial cells have reduced sensitivity to microbial exposure relative to the adult to prevent excessive immune reactivity (*61*). However, the microbiota induces a temporary protective mechanism in the neonatal oral epithelium by increasing salivary antimicrobial components that shape the oral bacterial composition and burden (*60, 61*). Here we show that neonatal oral *Candida* colonization and consequently IL-22-mediated mucosal remodeling prevents fungal epithelial invasion. Given that neonates mount a vigorous T cell response to microbial exposure rather than developing immunological memory (*62*), oral mucosal remodeling may play a key role to control colonizing fungi during early life. Understanding such early mechanisms are important for human health, because a deficiency of this remodeling process could translate into oral and systemic pathologies in adult life. Future studies will be required to determine epithelial remodeling and turnover to age-related differences in bystander and antigen-specific T cell activation to maintain mucosal homeostasis.

Collectively, our findings identify oral mucosal remodeling through non-canonical IL-22 receptor signaling as a critical determinant of resistance to mucosal fungal infection and highlight the importance of tissue-specific immune responses in the control of infectious disease.

## Acknowledgments

We thank Jay K. Kolls (Tulane University) for providing the *Il22ra1*^*E2a-cre*^ mice, Salomé LeibundGut-Landmann (University of Zurich) for the *Candida albicans* commensal-like strain CA101, James G. Rheinwald (Dana-Farber/Harvard Cancer Center) for providing the OKF6/TERT-2 cell line, and members of the Division of Infectious Diseases at Harbor-UCLA Medical Center for critical suggestions.

## Funding

NIH grant R01DE022600 (MS), R01AI177254 (SGF), R21AI159221, R56AI175328 (NJ), R37DE022550 (SLG), Division of Intramural Research of the NIAID (MSL), California Institute for Regenerative Medicine Stem Cell Biology Training Grant EDUC4-12837 (NM).

## Author contributions

Conceptualization: MS

Methodology: NM, JS, NS, AM, FA, AW, MSL, SLG, NJ, SGF, MS

Investigation: NM, JS, NS, AM, FA, AW, MS

Visualization: NM, JS, FA, AW, MS

Funding acquisition: NM, NJ, SLG, SGF, MS

Project administration: MS

Supervision: NJ, MS

Writing – original draft: MS

Writing – review & editing: NM, JS, NS, AM, FA, AW, MSL, SLG, NJ, SGF, MS

## Competing interests

The authors declare no competing interests.

## Data and materials availability

The authors declare that the data supporting the findings of this study are available within the paper and the accompanying supplementary information files. The high-throughput sequencing data from this study have been deposited with links to BioProject accession numbers xxx and xxx in the NCBI BioProject database (accession numbers will be provided once paper is accepted for publication).

## Notes

### Competing Interest Statement

The authors have declared no competing interest.

## References

1. N. M. Moutsopoulos, J. E. Konkel, Trends Immunol 39, 276 (Apr, 2018).

2. D. Zheng, T. Liwinski, E. Elinav, Cell Research 30, 492 (2020/06/01, 2020).

3. S. L. Gaffen, N. M. Moutsopoulos, Sci Immunol 5 (Jan 3, 2020).

4. M. Swidergall, S. LeibundGut-Landmann, Mucosal Immunol 15, 829 (May, 2022).

5. F. L. Mayer, D. Wilson, B. Hube, Virulence 4, 119 (Feb 15, 2013).

6. M. Swidergall, S. G. Filler, PLoS Pathog 13, e1006056 (Jan, 2017).

7. T. Y. Shao et al., Cell Host Microbe 25, 404 (Mar 13, 2019).

8. M. G. Netea et al., Science 352, aaf1098 (Apr 22, 2016).

9. M. G. Netea et al., Nature Reviews Immunology 20, 375 (2020/06/01, 2020).

10. I. D. Iliev, I. Leonardi, Nat Rev Immunol 17, 635 (Oct, 2017).

11. H. R. Conti et al., J Exp Med 206, 299 (Feb 16, 2009).

12. H. R. Conti et al., Cell Host Microbe 20, 606 (Nov 9, 2016).

13. M. Swidergall, N. V. Solis, M. S. Lionakis, S. G. Filler, Nat Microbiol 3, 53 (Jan, 2018).

14. M. Swidergall et al., Cell Rep 28, 423 (Jul 9, 2019).

15. N. Millet, N. V. Solis, M. Swidergall, Front Immunol 11, 555363 (2020).

16. F. A. Schonherr et al., Mucosal Immunol 8, 2 (2017).

17. F. R. Kirchner et al., Front Immunol 10, 330 (2019-February-27, 2019).

18. M. J. Bissell, A. Rizki, I. S. Mian, Current opinion in cell biology 15, 753 (2003).

19. R. Okumura, K. Takeda, Experimental & Molecular Medicine 49, e338 (2017/05/01, 2017).

20. T. J. Break et al., Science 371 (Jan 15, 2021).

21. T. Krausgruber et al., Nature 583, 296 (Jul, 2020).

22. L. C. Frazer, M. Good, Mucosal Immunol 15, 1181 (Jun, 2022).

23. N. Torow, T. W. Hand, M. W. Hornef, Immunity 56, 485 (Mar 14, 2023).

24. K. Agaronyan et al., Immunity 55, 895 (May 10, 2022).

25. A. K. Beppu et al., Nat Commun 14, 5814 (Sep 19, 2023).

26. C. L. Hayes et al., Scientific Reports 8, 14184 (2018/09/21, 2018).

27. J. A. Dudakov, A. M. Hanash, M. R. van den Brink, Annu Rev Immunol 33, 747 (2015).

28. N. V. Solis, S. G. Filler, Nat Protoc 7, 637 (Mar 8, 2012).

29. R. B. Presland, B. A. Dale, Crit Rev Oral Biol Med 11, 383 (2000).

30. R. B. Presland, R. J. Jurevic, J Dent Educ 66, 564 (Apr, 2002).

31. M. S. Lionakis, R. A. Drummond, T. M. Hohl, Nat Rev Immunol 23, 433 (Jul, 2023).

32. N. Hernández-Santos, S. L. Gaffen, Cell Host Microbe 11, 425 (May 17, 2012).

33. C. A. Lindemans et al., Nature 528, 560 (Dec 24, 2015).

34. J. R. Mock et al., Mucosal Immunol 7, 1440 (Nov, 2014).

35. J. Hur et al., Circulation 116, 1671 (Oct 9, 2007).

36. F. E. Y. Aggor et al., Sci Immunol 5 (Jun 5, 2020).

37. R. Bichele et al., Eur J Immunol 48, 464 (Mar, 2018).

38. E. Kaleviste et al., Front Immunol 11, 838 (2020).

39. M. W. Plank et al., J Immunol 198, 2182 (Mar 1, 2017).

40. M. Keir, Y. Yi, T. Lu, N. Ghilardi, J Exp Med 217, e20192195 (Mar 2, 2020).

41. S. Mossner et al., J Biol Chem 295, 12378 (Aug 28, 2020).

42. S. Rose-John, B. J. Jenkins, C. Garbers, J. M. Moll, J. Scheller, Nature Reviews Immunology 23, 666 (2023/10/01, 2023).

43. A. Puel et al., Curr Opin Allergy Clin Immunol 12, 616 (Dec, 2012).

44. S. C. Jameson, D. Masopust, Immunity 31, 859 (2009/12/18/, 2009).

45. M. W. Hornef, N. Torow, Immunology 159, 15 (Jan, 2020).

46. S. Patil, R. S. Rao, B. Majumdar, S. Anil, Front Microbiol 6, 1391 (2015).

47. H. R. Razzaghian et al., Front Immunol 12, 655027 (2021).

48. V. Oikonomou et al., N Engl J Med 390, 1873 (May 30, 2024).

49. M. Swidergall, Pathogens 8, 40 (Mar 25, 2019).

50. M. Schirmer et al., Cell 167, 1125 (Nov 3, 2016).

51. T. Arshad, F. Mansur, R. Palek, S. Manzoor, V. Liska, Frontiers in immunology 11, 2148 (2020).

52. C. B. Korol et al., J Clin Immunol 43, 406 (Feb, 2023).

53. V. Béziat et al., J Exp Med 217 (Jun 1, 2020).

54. T. Arlabosse et al., J Clin Immunol 43, 1566 (Oct, 2023).

55. Z. Qin et al., J Exp Med 221 (Mar 4, 2024).

56. P. Pandiyan et al., Immunity 34, 422 (Mar 25, 2011).

57. F. R. Kirchner, S. LeibundGut-Landmann, Mucosal Immunol (Jul 27, 2020).

58. A. H. Verma et al., Sci Immunol 2 (Nov 3, 2017).

59. K. Trautwein-Weidner et al., PLoS Pathog 11, e1005164 (Oct, 2015).

60. N. Koren et al., Cell Host Microbe 29, 197 (Feb 10, 2021).

61. K. Zubeidat, A. H. Hovav, Trends Immunol 42, 622 (Jul, 2021).

62. B. D. Rudd, Annu Rev Immunol 38, 229 (Apr 26, 2020).

