## Supplemental Material for "Non-canonical IL-22 receptor signaling remodels the mucosal barrier during fungal immunosurveillance"

**Affiliations**
[1]Division of Infectious Diseases, Harbor-UCLA Medical Center, Torrance, CA, USA
[2]The Lundquist Institute for Biomedical Innovation at Harbor-UCLA Medical Center, Torrance, CA, USA
[3]Division of Respiratory and Critical Care Medicine and Physiology, Harbor-UCLA Medical Center, Torrance, CA, USA
[4]University of Pittsburgh, Division of Rheumatology and Clinical Immunology, Pittsburgh, PA, USA
[5]Fungal Pathogenesis Section, Laboratory of Clinical Immunology and Microbiology (LCIM), National Institute of Allergy and Infectious Diseases (NIAID), Bethesda, MD, USA
[6]David Geffen School of Medicine at UCLA, Los Angeles, CA, USA

**Supplementary Material**

- Material & Methods
- Supplementary Figures (S1 – S21)
- References (63 - 72)

### •Material & Methods

**Ethics statement.** All animal work was approved by the Institutional Animal Care and Use Committee (IACUC) of the Lundquist Institute at Harbor-UCLA Medical Center and the University of Pittsburgh.

**Organisms and cell lines.** The *C. albicans* strains SC5314 (pathogenic-like; PL), and CA101 (commensal-like; CL) were used in the experiments and were grown as described previously (*15*). The OKF6/TERT-2 cells have been authenticated by RNA-sequencing (*12*), and have been tested for mycoplasma contamination. The intestinal epithelial cell line C2BBe1 was purchased from ATCC.

**Subject details.** For *in vivo* animal studies, age-and sex matched mice were used. Animals were bred and housed under pathogen-free conditions at the Lundquist Institute. Animals were randomly assigned to the different infection or treatment groups. Researchers were not blinded to the experimental groups because the endpoints (fungal burden, histology, cytokine levels) were objective measures of disease severity. *Il22ra1*$^{E2a\text{-}cre}$ (referred to as *Il22ra1*$^{-/-}$) mice were provided by Jay K. Kolls (*63*). *IL-22TdTomato* (*64*) mice were bred at the University of Pittsburgh. *Il10rb*$^{-/-}$ (B6.129S2-*Il10rb*$^{tm1Agt}$/J) (*65*), *Il22*$^{iCre/iCre}$ (C57BL/6-*Il22*$^{tm1.1(icre)Stck}$/J; referred to as *Il22*$^{-/-}$) (*66*), *Il10*$^{-/-}$ (B6.129P2-*Il10*$^{tm1Cgn}$/J) (*67*), and *Rag1*$^{-/-}$ (B6.129S7-*Rag1*$^{tm1Mom}$/J; non-leaky) (*68*) mice were purchased from the Jackson Laboratory, and maintained at the Lundquist Institute. C57BL/6 wild type mice were purchased from the Jackson Laboratory and cohoused with breeders for at least 1 week before the experiments.

**Mouse model of oropharyngeal candidiasis.** OPC was induced in mice as described previously (*15, 28*). Briefly, for inoculation, the animals (7-8 week old male and female mice) were sedated, and a swab saturated with $2 \times 10^7$ *C. albicans* cells was placed sublingually for 75 min. For colony-forming unit (CFU) enumeration the tongues were harvested, weighed, homogenized and quantitatively cultured. Neonatal mice were generated using standard breeding procedures and 8-week-old adult mice. For antibody depletion wild-type mice were treated intraperitoneally with 200 µg of anti-IL-22 (IL22JOP, Invitrogen) and 200µg of anti-mouse IL-17A (17F3, BioXCell), or corresponding concentrations of isotype controls (Rat IgG2a kappa, eBR2a, mouse IgG1 isotype, MOPC-21) on day 11 and 13 post oral CL-infection. For pharmacological gp130 inhibition, mice were treated daily starting one day prior CL colonization with 10mg/kg SC144 (SelleckChem) in veterinary 0.9% Sodium Chloride (Vetivex).

**Single cell RNA-sequencing.** Tongues single cell suspensions were generated as described previously (*14*). Briefly, mice were orally infected with *C. albicans* as described above. After 11 days of infection, the animals were administered a sublethal anesthetic mix intraperitoneally. The thorax was opened, and a part of the rib cage was removed to gain access to the heart. The vena cava was transected and the blood was flushed from the vasculature by slowly injecting 10 mL PBS into the right ventricle. The tongue was harvested and cut into small pieces in 100 µL of ice-cold PBS. 1 mL digestion mix (4.8 mg/ml Collagenase IV; Worthington Biochem, and 200 µg/ml DNase I; Roche Diagnostics, in 1x PBS) was added after which the tissue was incubated at 37°C for 45 min. The resulting tissue suspension was then passed through a 100 µm cell strainer. For one sample, single cells of two mice were combined. Single cell library preparation and sequencing were conducted by Singulomics (Bronx, USA). Cryopreserved viable single cell suspensions were thawed, washed, resuspended in cell culture media with 0.04% bovine serum albumin and counted. Viable cell suspensions were then loaded into the Chromium Controller (10x Genomics) to generate gel beads-in-emulsion (GEM), with each GEM containing a single cell as well as barcoded oligonucleotides. We targeted 10,000 cells to be captured per sample. Next, the GEMs were placed in the SimpliAmp 96-well Thermal Cycler (Thermo Fisher Scientific) and reverse transcription was performed in each GEM (GEM-RT). After the reaction, the complementary cDNA was amplified and cleaned using Silane DynaBeads (10x Genomics) and the SPRI select Reagent kit (Beckman Coulter). Amplified full-length cDNAs from poly-adenylated mRNA were then used to generate a 3′ Gene Expression library (Chromium Next GEM 3′ Single Cell Reagent kits v3.1, dual index) following the manufacturer's instructions (10x Genomics). Amplified cDNAs and the libraries were measured using Qubit dsDNA HS assay (Thermo Fisher Scientific) and quality was assessed using BioAnalyzer (Agilent Technologies). The libraries were sequenced with ~200 million PE150 reads per sample on Illumina NovaSeq 6000, and fastq files of two samples were generated using Illumina bcl2fastq. Fastq files were subsequently processed using 10x Genomics Cell Ranger analytical pipeline (v4.0.0) and mouse mm10 reference. Cellranger aggr was used to aggregate outputs from both the libraries and the aggregated filtered processed files were finally taken as input to do a quality control check and further downstream clustering analysis using Cellenics/Trailmaker software (Parse Biosciences). After quality control to remove low-quality cells expressing high mitochondrial gene signatures and exclude doublets, Louvain clustering identified 20 subgroups in our samples (Fig. S2). Cluster 1 and 10 were excluded from further analysis due to enriched lncRNA Gm42418 which is associated with ribosomal RNA contamination. The high-throughput sequencing data from this study have been submitted to the NCBI Sequence Read Archive (SRA) under accession number xxx (number will be provided once paper is accepted for publication).

**Epithelial- enriched RNA-sequencing.** The tongue tissue was placed in a 60mm petri dish, and epithelial cells were isolated by gentle mechanical isolation. The isolated cells were placed in lysis buffer reagent and homogenized in Lysing Matrix C tubes (MPBio) containing a ¼” ceramic bead (MPBio). RNA extraction was performed with the RiboPure Kit (Ambion), and RNA sequencing was performed by Novogene Corporation Inc. (Sacramento, USA) as described previously (*69*). mRNA was purified from total RNA using poly-T oligo-attached magnetic beads. To generate the cDNA library the first cDNA strand was synthesized using random hexamer primer and M-MuLV Reverse Transcriptase (RNase $H^-$). Second strand cDNA synthesis was subsequently performed using DNA Polymerase I and RNase H. Double-stranded cDNA was purified using AMPure XP beads and remaining overhangs of the purified double-stranded cDNA were converted into blunt ends via exonuclease/polymerase. After 3' end adenylation a NEBNext Adaptor with hairpin loop structure was ligated to prepare for hybridization. In order to select cDNA fragments of 150~200 bp in length, the library fragments were purified with the AMPure XP system (Beckman Coulter, Beverly, USA). Finally, PCR amplification was performed and PCR products were purified using AMPure XP beads. The samples were read on an Illumina NovaSeq 6000 with ≥20 million read pair per sample. Downstream analysis was performed using a combination of programs including STAR, HTseq, and Cufflink. Alignments were parsed using Tophat and differential expressions were determined through DESeq2. KEGG enrichment was implemented by the ClusterProfiler. Gene fusion and difference of alternative splicing event were detected by Star-fusion and rMATS. The reference genome of *Mus musculus* (GRCm38/mm10) and gene model annotation files were downloaded from NCBI/UCSC/Ensembl. Indexes of the reference genome was built using STAR and paired-end clean reads were aligned to the reference genome using STAR (v2.5). HTSeq v0.6.1 was used to count the read numbers mapped of each gene. The FPKM of each gene was calculated based on the length of the gene and reads count mapped to it. FPKM, Reads Per Kilobase of exon model per Million mapped reads, considers the effect of sequencing depth and gene length for the reads count at the same time (*70*). Differential expression analysis was performed using the DESeq2 R package (2_1.6.3). The resulting P-values were adjusted using the Benjamini and Hochberg’s approach for controlling the False Discovery Rate (FDR). Genes with an adjusted P-value <0.05 found by DESeq2 were assigned as differentially expressed (cutoff fold change 1.5, table S1).To allow for log adjustment, genes with 0 FPKM are assigned a value of 0.001. Correlation were determined using the cor.test function in R with options set alternative =”greater” and method = “Spearman”. To identify the correlation between the differences, we clustered different samples using expression level FPKM to see the correlation using hierarchical clustering distance method with the function of heatmap, SOM (Self-organization mapping) and kmeans using silhouette coefficient to adapt the optimal classification with default parameter in R. We used clusterProfiler R package to test the statistical enrichment of differential expression genes in KEGG pathways. The high-throughput sequencing data from this study have been submitted to the NCBI Sequence Read Archive (SRA) under accession number xxxx (will be provided when manuscript accepted for puplication).

**Immunofluorescence.** To determine host cell localization *in vivo*, 15-30-μm-thick sections of OCT-embedded tongues were fixed with cold acetone. Next, the cryosections were rehydrated in PBS and then blocked using BSA. To detect K14 (#Ab181595, Abcam), K13 (#BSM-52053R, Bioss), Ki67(#9126S, Cell Signaling), or CD4 (#100405, biolegend) positive cells, the sections were incubated with a primary antibody overnight followed by 1h incubation at room temperature with a secondary antibody either Alexa Fluor 488 or 568 goat anti- rabbit IgG (A11034, #A11011). To detect *C. albicans*, the sections were also stained with an anti-*Candida* antiserum (Biodesign International) conjugated with Alexa Fluor 568 (Thermo Fisher Scientific) for 1 hour. To visualize the nuclei, the cells were also stained with DAPI (4′,6-diamidino-2-phenylindole).The sections (z-stack) were imaged by confocal microscopy. To enable comparisons of fluorescence intensities among slides, the same image acquisition settings were used for each experiment.

**Histology**. Half-tongues were embedded in OCT (Fisher HealthCare, Houston, USA), frozen on dry ice, and stored at -80°C. Sagittal cryosections (5 μm) were air-dried at room temperature, stained with periodic acid-Schiff (Sigma-Aldrich), and counterstained with hematoxylin (Sigma-Aldrich). Sections were mounted with a toluene solution (Permount®; Fisher Chemical). Images were analyzed with a phase-contrast microscope (Zeiss Axiostar) and Gryphax Software.

**Cytokine and chemokine measurements *in vivo*.** To determine the whole tongue cytokine and chemokine protein concentrations, mice were infected as described above. The mice were euthanized at various time points, and their tongues were harvested, weighed, and homogenized. The homogenates were cleared by centrifugation and the concentration of inflammatory mediators was measured using the Luminex multiplex bead assay (Invitrogen). In some experiments, IL-17A and IL-22 (R&D Systems) were determined by ELISA according to the manufacturer's instructions.

**Immunophenotyping.** Mice were orally infected with *C. albicans* as described above. After different time points, the animals were administered a sublethal anesthetic mix intraperitoneally. The thorax was opened, and a part of the rib cage removed to gain access to the heart. The vena cava was transected and the blood was flushed from the vasculature by slowly injecting 10 ml PBS into the right ventricle. The tongue was harvested and cut into small pieces in 100 μl of ice-cold PBS. 1 ml digestion mix (4.8 mg/ml Collagenase IV; Worthington Biochem, and 200 μg/ml DNase I; Roche Diagnostics, in 1x PBS) was added after which the tissue was incubated at 37°C for 30 min. The resulting tissue suspension was then passed through a 100 μm cell strainer. Cell suspensions were separated by Percoll gradient centrifugation as described before (*15, 36*). To determine IL-17A and IL-22, cell suspensions were stimulated with Pharmingen™ Leukocyte Activation Cocktail (BD Biosciences) for 5 hours. Cell were washed and stained with CD4 antibody (company). For intracellular staining, cells were fixed with Cytofix/Cytoperm (BD Biosciences) and stained for 1 hour with IL-17A (company) and IL-22(company) antibodies. The stained cells were analyzed on FACSymphony system (BD Biosciences), and the data were analyzed using FACS Diva and FlowJo software. Th17 cells were identified as singlets CD4+ IL-17A+ IL22+ and Th22 cells were identified as singlets CD4+ IL-17A- IL22+ (Fig. S19).

**ELISpot.** Cervical lymph nodes (cLN) were isolated from CL-colonized or Sham-infected animals. cLN were digested with 2.4 mg/ml Collagenase I (Thermo Fisher) and 200 μg/ml DNase I (Roche Diagnostics) in 1x PBS. The resulting tissue suspension was then passed through a 70 μm cell strainer, washed, and live cells were determined. 1x$10^5$ cLN cells were plated on pre-coated PVDF plate, left unstimulated or were stimulated with 15 µg *Candida* peptide pool for 24 hrs at 37°C and 5% $CO_2$. ELISpot was performed according to the manufacturer's instructions (ImmunoSpot). Plates were analyzed with a ImmunoSpot® Analyzer (CTL). For *Candida*- specific IL-17 detection, spots of *Candida* pool wells were determined and subtracted from spots determined in corresponding unstimulated wells. The peptide pool (lysate) was generated from stationary cultures of CL strain CA101 as described elsewhere (*71*).

**Live-cell imaging for cell proliferation.** 1.2×$10^4$ OKF6/TERT-2 cells were plated in 96-well plates and incubated overnight at 37ºC, allowing them to settle down. Cells were treated without or with 10, 25, 50, 100, and 200 ng/ml of human IL-22 (PeproTech) in keratinocyte serum-free medium (KSF), and immediately placed in the IncuCyte SX5 Live Cell Analysis System (Sartorius, Göttingen, Germany). To identify the impacts of IL-22-mediated receptor complex and downstream signaling pathways, cells were treated with 5µg/ml anti-hIL-22Rα1 (AF2770; R&D Systems), 5µg/ml anti-IL-10RB (90220; R&D Systems) or in a combination, 1µM TYK2 inhibitor (BMS-911543, SelleckChem), 0.5µM (C188-9; SelleckChem), 5µM STAT3 inhibitor (S3I-201; SelleckChem), 50 nM gp130 inhibitor (SC144; SelleckChem), and 5µg/ml tocilizumab (A2012; SelleckChem). Cell proliferation was observed and automatically analyzed from acquired three images per well taken every four hours for a consecutive 72 hours. Data was analyzed with the Incucyte live cell analysis software.

**Immunoblotting.** OKF6/TERT-2 cells in 24-well tissue culture plates were switched to KSF medium without supplements for 1 h and treated with 50 ng/mL recombinant human IL-22 (PeproTech). Next, the cells were rinsed with cold HBSS containing protease and phosphatase inhibitors and detached from the plate with a cell scraper. After the cells were collected by centrifugation, they were boiled in sample buffer. The lysates were separated by SDS-PAGE, and phosphorylation and total proteins were detected by immunoblotting with specific antibodies against pSTAT3 (Tyr705, D3A7, #9145, Cell Signaling), STAT3 (D3Z2G, #12640, Cell Signaling), and β-actin (8H10D10, #3700, Cell Signaling). Each experiment was performed at least 3 times. The immunoblots were developed using enhanced chemiluminescence and imaged with a C400 digital imager (Azure Biosystems, Dublin, CA, USA). Uncropped immunoblots are presented in Fig. S20-21.

**Co-immunoprecipitation.** OKF6/TERT-2 cells were grown in KSF medium with supplements to 70% confluence and cells were treated without or with 50ng/ml human IL-22 (PeproTech). After 3 hrs. cells were incubated with fresh media containing 0.1mM DSP-crosslinking (dithiobis(succinimidyl propionate) for 30min. Cells were washed twice with PBS and re-incubated with fresh media containing 25mM Tris-HCl for 15min. Cells were lysed with ice-cold IP lysis buffer with protease/phosphatase inhibitors and protein concentrations were measured by Pierce BCA protein assay (Thermo Scientific). Co-immunoprecipitation was performed using the Pierce Classic Magnetic IP/Co-IP Kit according to the manufacturer's protocol (Thermo Scientific). In brief, 400µg protein samples were added with 15µg IL22RA1 (AF2770; R&D systems), IL10RB (MAB874; R&D systems), or gp130 antibody (362002; BioLegend). The protein-antibody mixture was incubated overnight on a rotator at 4 ºC to facilitate immune-complex formation. Following overnight incubation, 25 µL pre-washed Pierce Protein A/G magnetic beads were added to the protein-antibody mixture to capture the immuno-complex. The mixture was incubated for 1h on a rotator at room temperature. Proteins were collected using a magnetic stand and separated by SDS-PAGE.

**Proximity ligation assay.** 2.5x$10^5$ OKF6/TERT-2 cells were seeded onto fibronectin-treated coverslips in a 24-well plate overnight. The next day, cells were fixed with 2% paraformaldehyde for 10 minutes at room temperature. After washing and blocking, the cells were incubated with a goat anti-IL22RA antibody (#AF2770, R&D Systems, 5µg/ml), a mouse anti-IL10RB antibody (#MAB874, Clone 90220, R&D Systems, 5µg/ml), a rabbit anti-gp130 antibody (#37325, Cell

Signaling, 5μg/ml), or two of these antibodies in combination overnight at 4°C. The interactions of IL22RA with IL10RB and the interactions of gp130 with either IL22RA or IL10RB were detected using the Duolink In Situ Red Starter Kit (DUO92101-1kit, Sigma-Aldrich) following manufacturer's instructions. The cells were imaged using a Leica TCS SP8 confocal microscope. Z-stacked images were obtained and overlaid using the Leica image processing software. To calculate the average number of complexes per cell, images were analyzed using ImageJ (*72*).

**Quantification and statistical analysis.** At least three biological replicates were performed for all *in vitro* experiments unless otherwise indicated. Data were compared by Ordinary one-way ANOVA or Mann–Whitney corrected for multiple comparisons were appropriate using GraphPad Prism (V. 10.3) software. P values <0.05 were considered statistically significant.

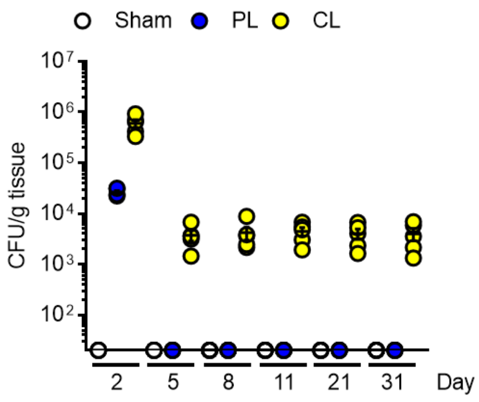


**Figure S1 The commensal-like (CL) isolate CA101 persists in the oral cavity**. Oral fungal burden of wild-type mice infected with indicated strains. Results are median ± standard error of the mean (SEM) of four independent experiments (*N* = 5/group). The y-axis is set at the limit of detection (20 CFU/g tissue). PL, pathogen-like; CL, commensal-like.

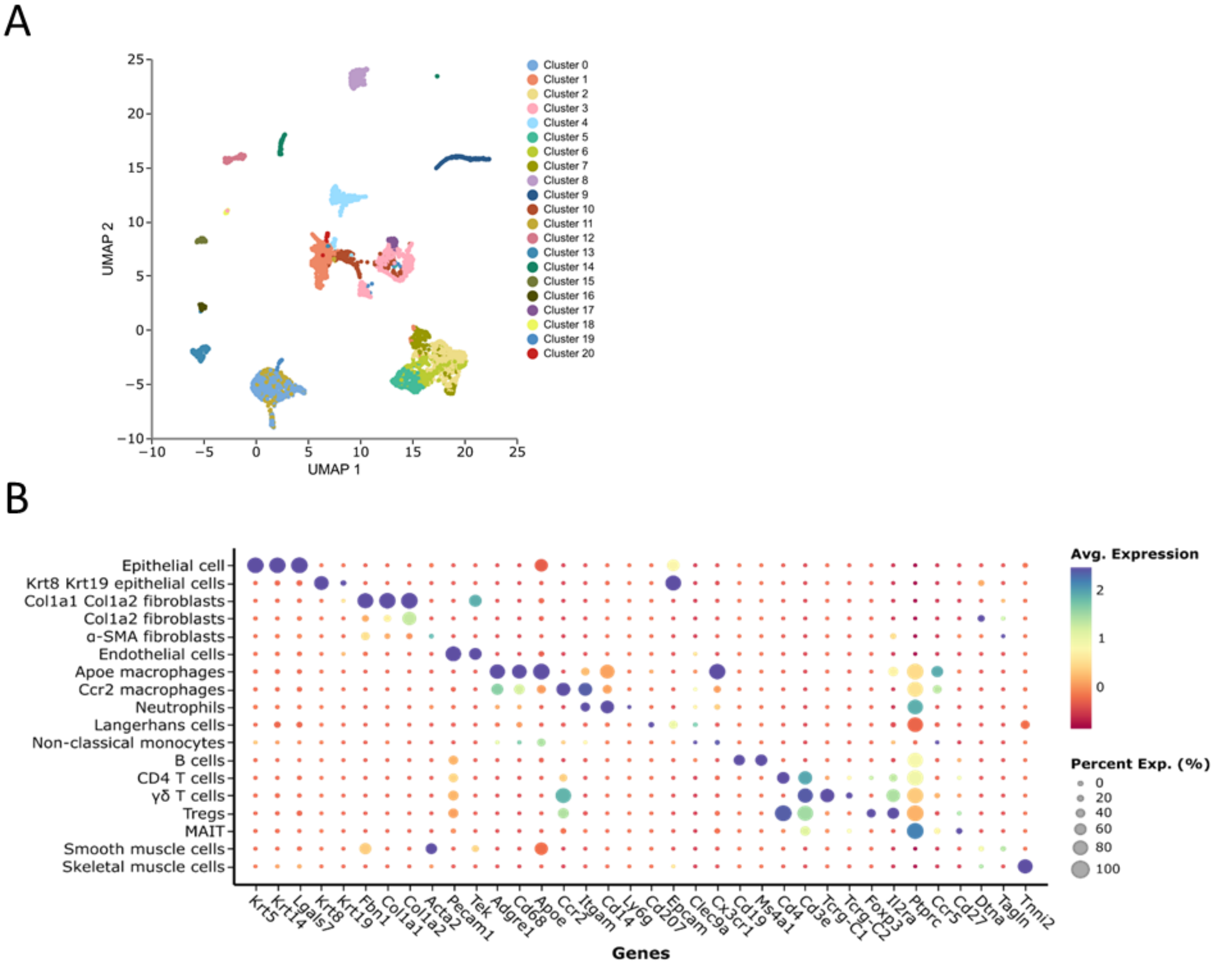


**Figure S2 Analysis of mucosal single cell RNA sequencing. A** Louvain clustering of single cells isolated from the tongue of CL- and PL-infected mice after 11 days of infection. UMAP, Uniform Manifold Approximation and Projection for Dimension Reduction. **B** Dot plot of scaled cell-type specific marker gene expression of identified subpopulations.

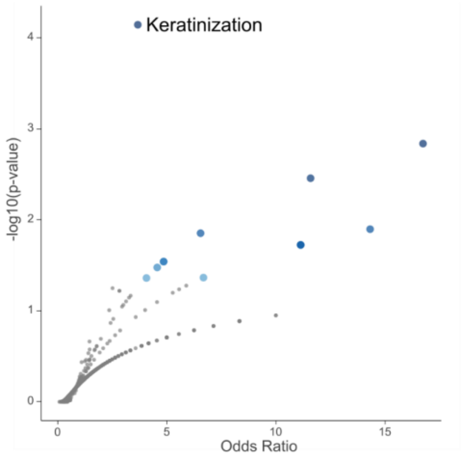


**Figure S3 Enriched pathways in epithelial cells from colonized mice in the scRNA-sequencing data set.** Upregulated genes identified in the epithelial cell subset were analyzed for pathway enrichment. –log 10 p value is plotted against the odds ratio.

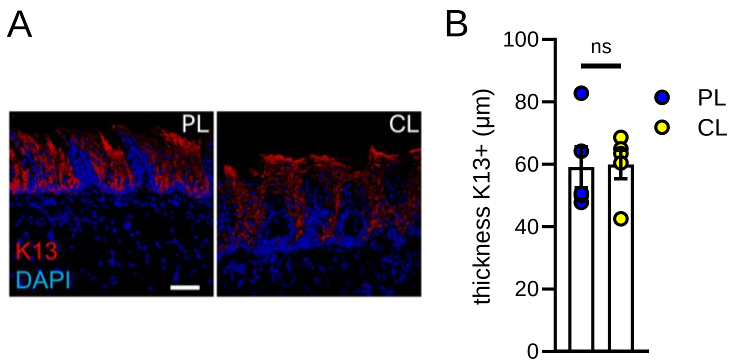


**Figure S4 K13 distribution remains the same during CL-colonization.** (A) Representative immunofluorescence pictures of keratin 13 (K13) after 11 days of infection. Mice were infected with PL and CL. Scale bar 50µm. (B) K13 thickness 11 days post infection. *N* = 5. Two-tailed Mann-Whitney

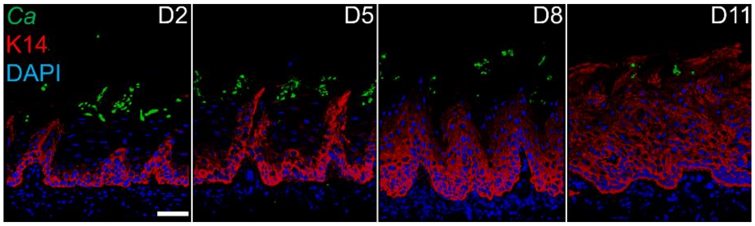


**Figure S5 Time course studies of K14 epithelial expansion.** Representative immunofluorescence pictures of keratin 14 (K14) after indicated time points. Mice were infected with CL. Scale bar 50μm.

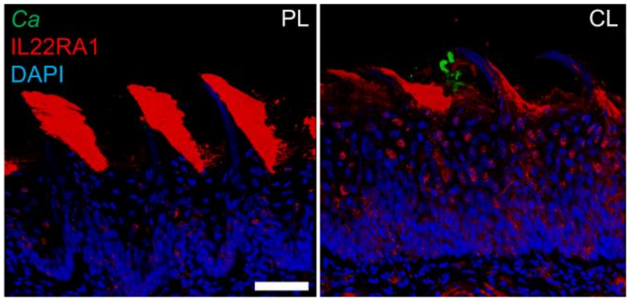


**Figure S6 IL-22RA1 distribution during fungal colonization in the oral cavity.** Representative immunofluorescence pictures Pof IL-22RA1 after 11 days of infection. Mice were infected with PL and CL. Scale bar 50μm.

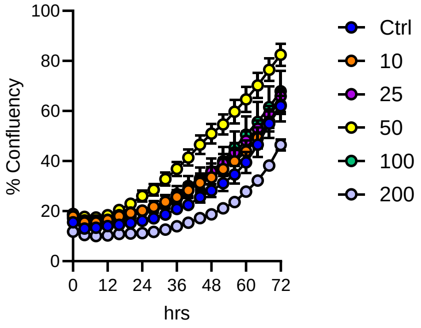


**Figure S7 Effect of different doses of IL-22 on oral epithelial cell proliferation.** Growth curve of human oral epithelial cells in the presence of different amounts recombinant IL-22. *N*=8.

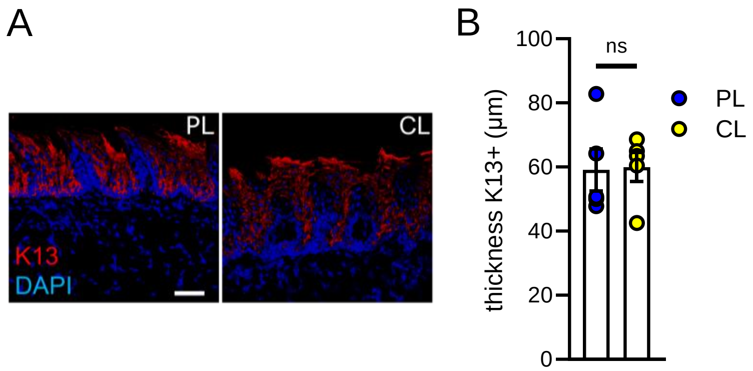


**Figure S8 Effect of IL-22 depletion on K14 expansion during oral fungal colonization** (A) Representative immunofluorescence pictures of keratin 14 (K14) after 11 days of infection. Mice were infected with CL and treated with isotype or α-IL-22 antibody starting day -1 every other day. Scale bar 50µm.

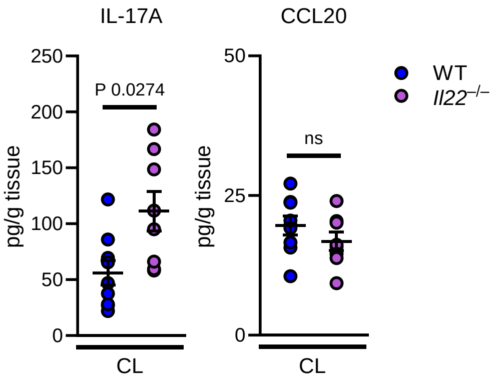


**Figure S9 Intact IL-17 signaling in *Il22*$^{-/-}$ mice during Candida colonization.** Levels of IL-17A and CCL20 in tongue homogenates after 11 days of CL colonization. Combined data of two independent experiments. N=8-9. Two-tailed Mann-Whitney.

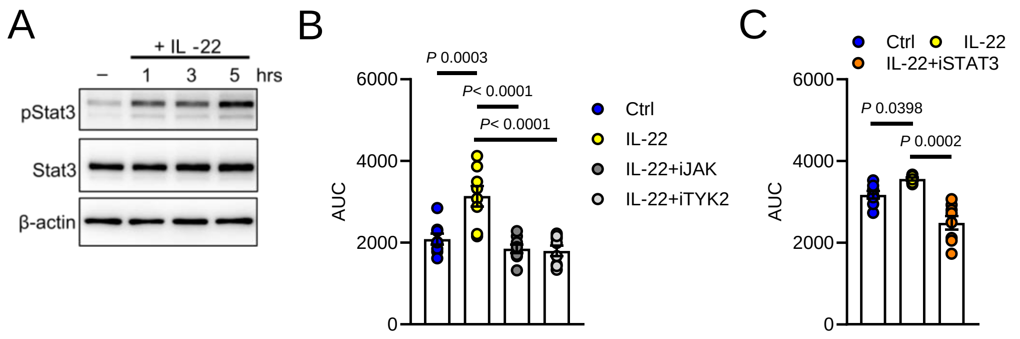


**Figure S10 JAK-STAT3 signaling is required for oral epithelial proliferation. A** Representative immunoblot of STAT3 activation during IL-22 incubation with 50ng/ml. **B** Area under the curve (AUC) of oral epithelial cells in response to IL-22 treatment in the presence of JAK inhibitor (iJak) and TYK2 inhibitor (iTyk2). Growth was determined by confluence over time. *N* = 8. Ordinary one-way ANOVA. **C** Area under the curve (AUC) of oral epithelial cells in response to IL-22 treatment in the presence of STAT3 inhibitor (iSTAT3) and Tyk2 inhibitor (iTYK2). Growth was determined by confluence over time. *N* = 8. Ordinary one-way ANOVA.

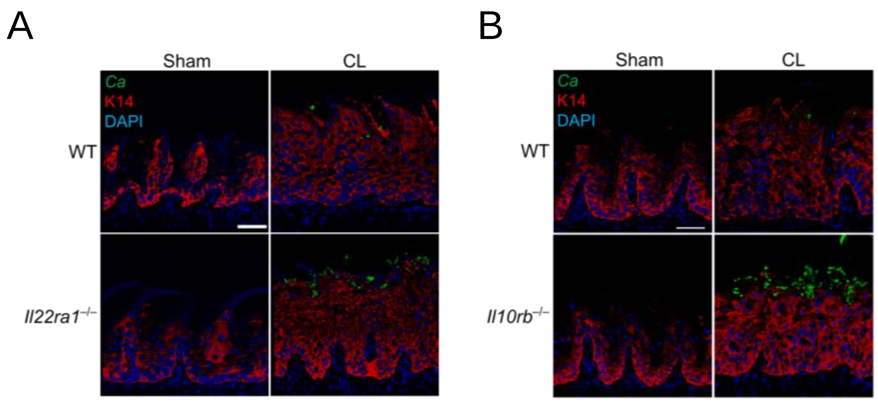


**Figure S11 K14 epithelial expansion during CL-colonization in *Il22ra1*$^{-/-}$ and *Il10rb*$^{-/-}$ mice.** Representative immunofluorescence pictures of K14 after 11 days of Sham- or CL infection of *Il22ra1*$^{-/-}$ (**A**) and *Il10rb*$^{-/-}$ (**B**) mice. Mice were infected with CL. Scale bar 50μm. Note pictures

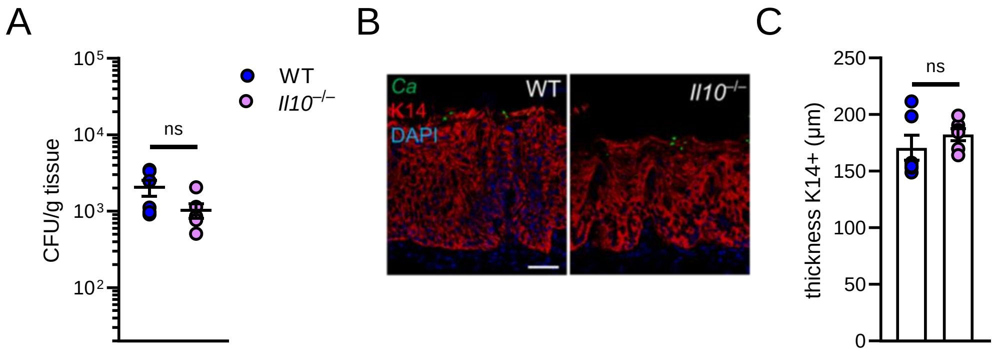


**Figure S12 IL-10 is dispensable to control oral fungal burden and K14 expansion during CL-colonization.** (A) Oral fungal burden of WT and *Il10*$^{-/-}$ mice colonized with CL after 11 days. *N* = 6; combined data of two independent experiments. Two-tailed Mann–Whitney Test. **B** Representative immunofluorescence pictures of K14 after 11 days of infection of indicated mice. Mice were infected with CL. Scale bar 50 µm. **C** Quantification of K14 thickness in indicated mice at day 11. *N* = 6; combined data of two independent experiments. Mann–Whitney Test.

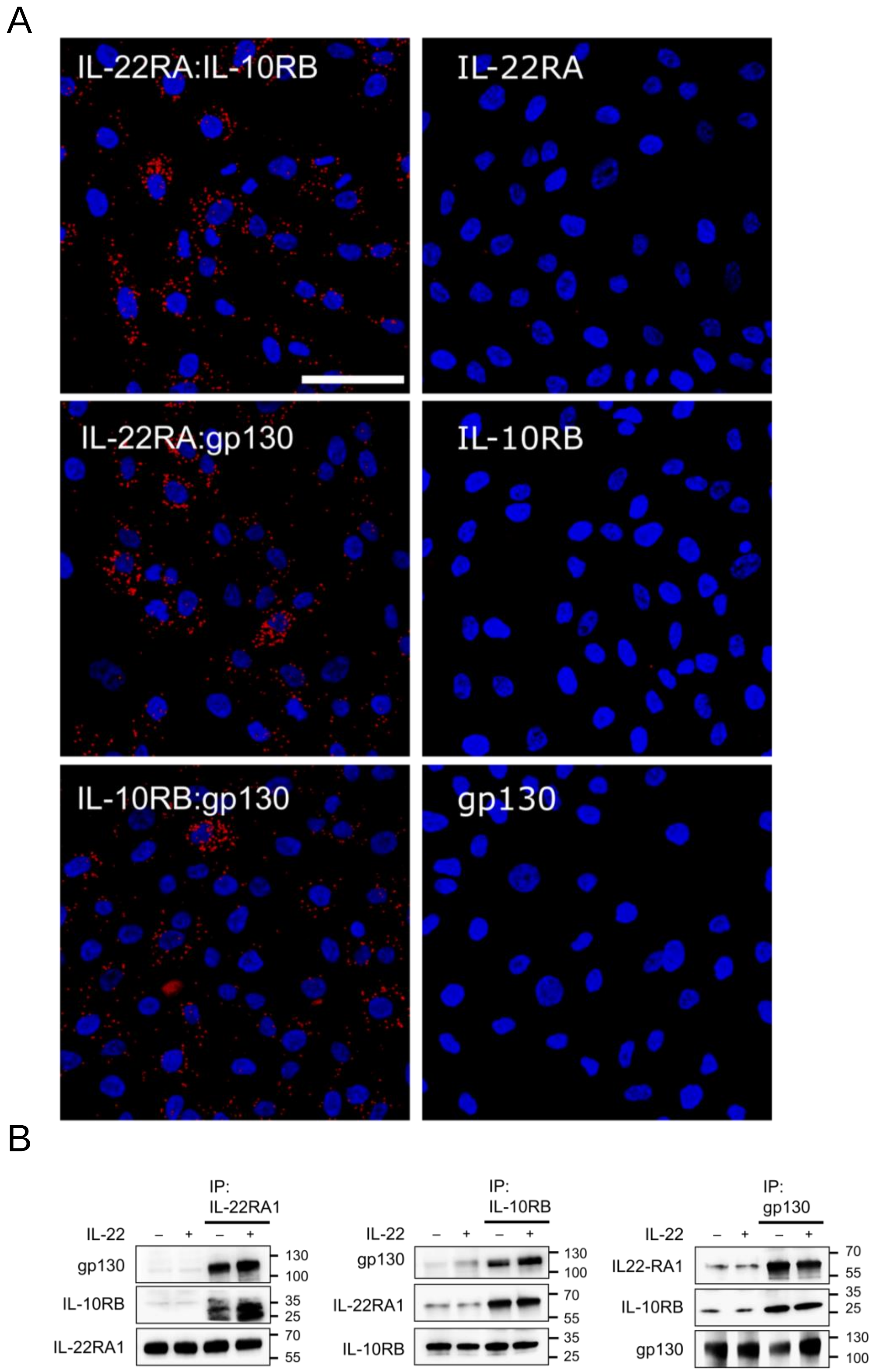
A
IL-22RA:IL-10RB
IL-22RA
IL-22RA:gp130
IL-10RB
IL-10RB:gp130
gp130
B
IP: IL-22RA1
IP: IL-10RB
IP: gp130
IL-22
– + – +
gp130
IL-10RB
IL-22RA1
IL22-RA1
130
100
70
55
35
25

**Figure S13 gp130 forms receptor complexes with IL-10RB and IL-22RA1. A** Representative images for proximity ligation assay (PLA) for indicated receptor complexes. Single antibody incubation was used as control Scale bar 50µm. Red dots indicate receptor complexes. **B** Representative immunoblot of immunoprecipitation of receptor complexes. OKF6-TERT2 cells were stimulated for 180 min with 50 ng/ml IL-22. After lysis, proteins (IL-22RA1, IL-10RB, and gp130) were immune-precipitated (IP) using specific antibodies. Lysates and pull-down samples (IP) were analyzed by immunoblotting. $N = 3$.

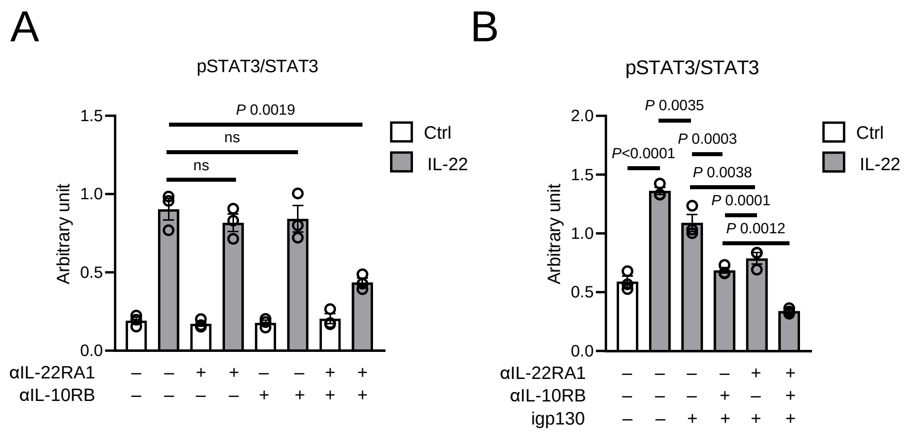


**Figure S13 Quantification of STAT3 activity during IL-22 incubation. A** Densitometric analysis (arbitrary units) of STAT3 phosphorylation normalized to total STAT3 protein in the presence of IL-22RA1, IL-10RB, or combination of IL-22RA1/IL-10RB blocking antibodies. *N* = 3. Ordinary one-way ANOVA. **B** Densitometric analysis (arbitrary units) of STAT3 phosphorylation normalized to total STAT3 protein in the presence of IL-22RA1, IL-10RB, igp130 (gp130 inhibitor), or combination. *N* = 3. Ordinary one-way ANOVA.

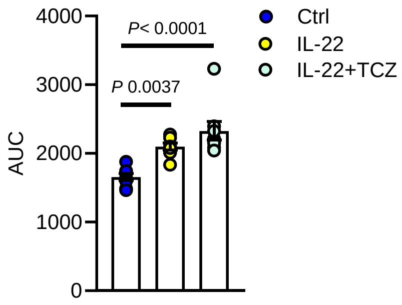


**Figure S15 Oral epithelial cell proliferation in the presence of tocilizumab.** Area under the curve (AUC) of oral epithelial cells in response to IL-22 treatment in the presence of tocilizumab. Growth was determined by confluence over time. *N* = 8. Ordinary one-way ANOVA.

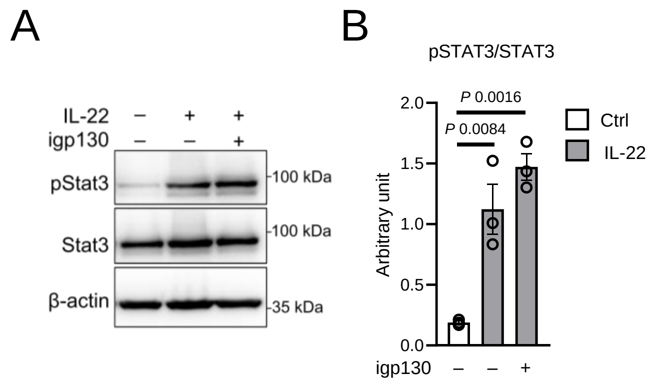


**Figures S16 STAT3 activation in intestinal epithelial cells is independent of gp130 signaling. A** Representative immunoblot of STAT3 activation in intestinal epithelial cells during IL-22 incubation in the presence of gp130 inhibitor. **B** Densitometric analysis (arbitrary units) of STAT3 phosphorylation normalized to total STAT3 protein in the presence of igp130 (gp130 inhibitor) in intestinal epithelial cells. *N* = 3. Ordinary one-way ANOVA.

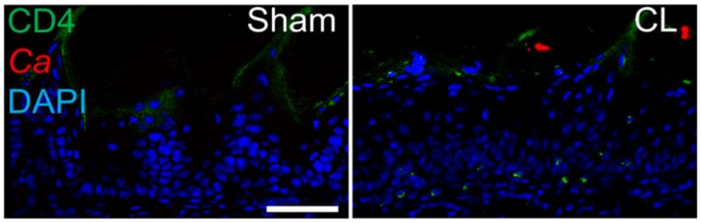


**Figures S17 CD4 T cell infiltration at the onset of neonatal colonization.** Representative immunofluorescence pictures of CD4 cells after 12 days of CL colonization of 2-day-old neonates mice. Scale bar 50μm.

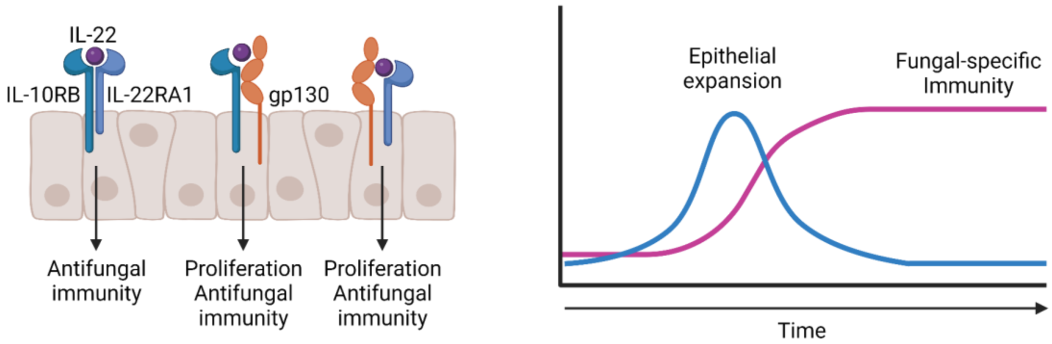


**Figure S18 Commensal fungi remodel the mucosal epithelial barrier.** Left: IL-22 responsive receptor complexes promoting antifungal immunity and epithelial remodeling. Right: Fungal colonization generates a primary epithelial remodeling response followed by the generation of fungal-specific immunity and auxiliary epithelial remodeling. IL-22, interleukin-22; IL-10RB, interleukin 10 receptor subunit beta; Il-22RA1, interleukin 22 receptor subunit alpha 1; gp130, glycoprotein 130.

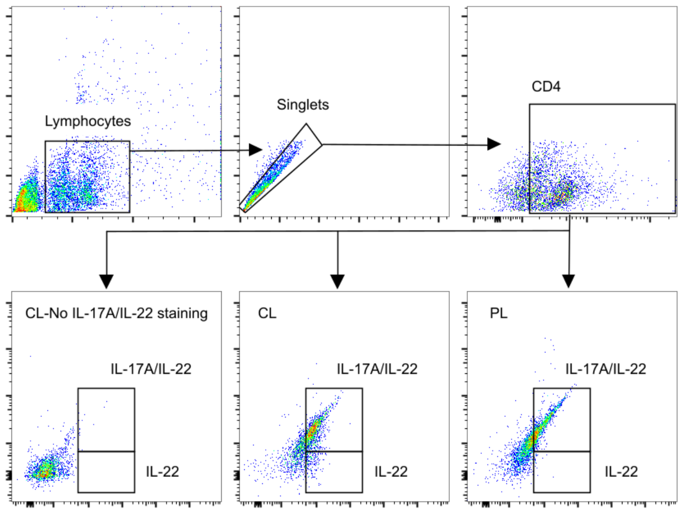


**Figure S19 Gating strategy of Th17/Th22 cells in the oral mucosa.** Th17 cells were identified as singlets CD4+ IL-17A+ IL22+ and Th22 cells were identified as singlets CD4+ IL-17A- IL22+.

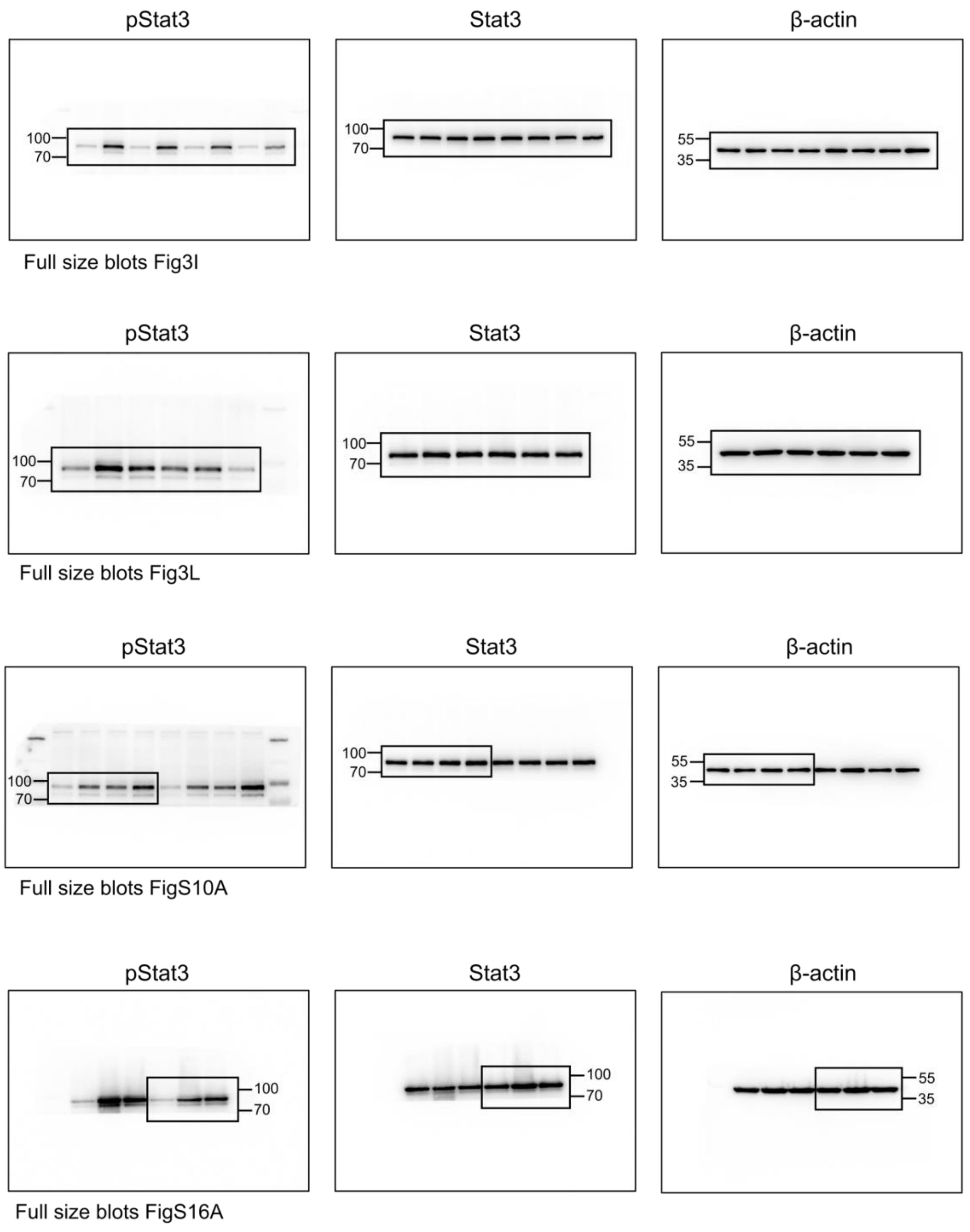


**Figure S20 Uncropped immunoblots.** Uncropped immunoblots of indicated figures.

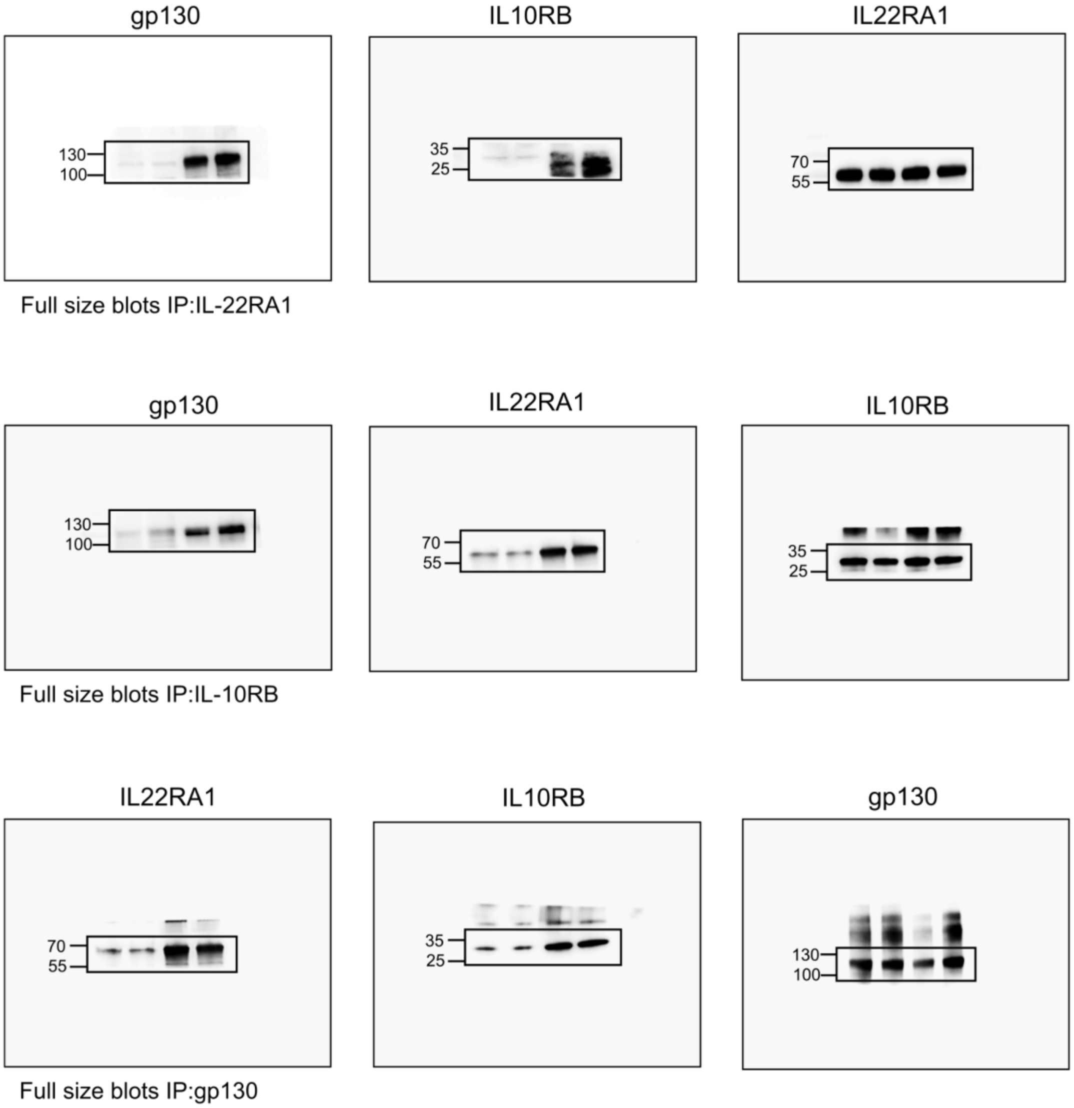


**Figure S21 Uncropped immunoblots of immunoprecipitation.** Uncropped immunoblots of indicated IPs.

### References


63. N. Iwanaga *et al.*, *Sci Immunol* **6**, eabf1198 (Sep 10, 2021).
64. W. Shen, J. A. Hixon, M. H. McLean, W. Q. Li, S. K. Durum, *Front Immunol* **6**, 662 (2015).
65. S. D. Spencer *et al.*, *J Exp Med* **187**, 571 (Feb 16, 1998).
66. H. Ahlfors *et al.*, *J Immunol* **193**, 4602 (Nov 1, 2014).
67. R. Kühn, J. Löhler, D. Rennick, K. Rajewsky, W. Müller, *Cell* **75**, 263 (Oct 22, 1993).
68. P. Mombaerts *et al.*, *Cell* **68**, 869 (Mar 6, 1992).
69. N. Millet *et al.*, *Nat Commun* **13**, 5545 (Sep 22, 2022).
70. A. Mortazavi, B. A. Williams, K. McCue, L. Schaeffer, B. Wold, *Nat Methods* **5**, 621 (Jul, 2008).
71. E. Szafranski-Schneider *et al.*, *PLoS Pathog* **8**, e1002501 (Feb, 2012).
72. C. T. Rueden *et al.*, *BMC Bioinformatics* **18**, 529 (Nov 29, 2017).
